# Long noncoding RNA *AVAN* promotes antiviral innate immunity by interacting with TRIM25 and enhancing the transcription of FOXO3a

**DOI:** 10.1101/623132

**Authors:** Chengcai Lai, Lihui Liu, Qinghua Liu, Sijie Cheng, Keyu Wang, Lingna Zhao, Min Xia, Cheng Wang, Hongjing Gu, Yueqiang Duan, Zhongpeng Zhao, Lili Zhang, Ziyang Liu, Jianjun Luo, Jianxun Song, Penghui Yang, Runsheng Chen, Xiliang Wang

## Abstract

Accumulating evidence has shown that long noncoding RNAs (lncRNAs) are involved in several biological processes, including immune responses. However, the role of lncRNAs in antiviral innate immune responses remains largely unexplored. Here, we identify an uncharacterized human lncRNA from influenza A virus (IAV) patients, antivirus and activate neutrophil (*AVAN*), that is significantly up-regulated upon virus infection. Mechanistically, nuclear lncRNA-*AVAN* positively regulates the transcription of forkhead box O3A (FOXO3a) by associating with its promoter and inducing chromatin remodeling to promote neutrophil chemotaxis. Furthermore, we also found that cytoplasmic lncRNA-*AVAN* directly binds tripartite motif containing 25 (TRIM25) and enhances the association of TRIM25 and Retinoic acid inducible gene-1 proteins (RIG-I) and the ubiquitylation of RIG-I, thereby promoting TRIM25- and RIG-I-mediated antiviral innate immune signaling. More importantly, we enforced the expression of AVAN in transgenic mice and found that it significantly alleviated IAV virulence and virus production. Collectively, these findings highlight the potential clinical implications of lncRNA-*AVAN* as a key positive regulator of the antiviral innate immune response and a promising target for developing broad antiviral therapeutics.

## Introduction

Long noncoding RNAs (lncRNAs), a large class of noncoding RNAs with no or limited coding potential, are defined as functional RNAs that participate in a wide range of biological processes, including cell and organ development, X-chromosome inactivation, tumor proliferation, genomic imprinting, and stem cell self-renewal (1–10). lncRNAs are also reportedly involved in innate and adaptive immune responses, such as immune cell proliferation and cytokine production (11–14). Previous studies have shown that lncRNAs exert their roles in regulating chromatin accessibility, mRNA stability and protein activity by interacting with chromatin DNA, mRNAs or proteins (15–18). With the discovery and characterization of an increasing number of lncRNAs, several mechanisms of these lncRNAs in virus-host interactions have been elucidated (19, 20), however, many others remain uncharacterized.

Influenza A virus (IAV) is a leading cause of respiratory-related morbidity and mortality, posing a substantial threat to global health (21). However, some mechanisms underlying IAV-host interactions remain unclear. Neutrophils provide one of the early lines of innate immunity and defense against invading microorganisms, which contribute to the fine regulation of the inflammatory and antiviral immune responses (22–24). Increasing evidence suggests that neutrophils are also prominent components of the inflammatory and immune responses during virus infection. Neutrophils are rapidly recruited to sites of infection during the innate immune response to IAV (25–27). In addition, neutrophils can produce and release a large variety of cytokines and chemokines (23), which enables them to significantly influence antiviral defense (28). During virus infection, chemokines play a crucial role in neutrophil activation and chemotaxis (29). However, how lncRNA functions in this process is largely unknown.

Intracellular RIG-I-like receptors (RLRs), namely, RIG-I, MDA5, and LGP2, are well-defined single-stranded viral RNA sensors that play a pivotal role in host antiviral activity by initializing the rapid production of type I interferons (IFN) and cytokines (30). RIG-I is a key sensor of paramyxoviruses, influenza virus, hepatitis C virus, and Japanese encephalitis virus (31). The binding of viral RNA to the RIG-I C-terminal regulatory domain results in a conformational change that, in turn, enables RIG-I binding to the signal adaptor MAVS (also called VISA, IPS-1, or CARDIF) through N-terminal caspase recruitment domains (CARDs). Such binding enhances the phosphorylation of IRF3 and eventually leads to the production of type I IFN and inflammatory cytokines (32). However, little is known about the roles of lncRNAs in IAV-infected patients or the mechanisms of lncRNA activity and host antiviral immune responses.

In this study, we profile the lncRNAs of IAV-infected patient neutrophils and identify a human lncRNA, designated *AVAN*, which plays a critical role in anti-IAV infection. Functional experiments demonstrate that *AVAN* up-regulates FOXO3a expression to promote neutrophil chemotaxis and recruitment. In addition, *AVAN* enhances the activation of the RIG-I-mediated antiviral response by directly binding to TRIM25. These results show that an unannotated lncRNA, *AVAN*, functions to maintain innate immune homeostasis of antiviral immunity during IAV infection.

## Results

### LncRNA expression is regulated by the influenza A (H7N9) virus in human neutrophils

Neutrophils, the prototypic cells of the innate immune system, are reported to play a pivotal role in the innate immune response to IAV (22–24). To explore the roles of host lncRNAs during IAV infection, we profiled whole transcriptional alterations using RNA-Seq in neutrophil samples from patients infected with IAV in the acute stage and their matched recovery-stage samples (GSE108807). We identified a total of 404 differentially expressed protein-coding genes (FC>2, *p<0.05*), including 234 up-regulated and 170 down-regulated genes, in each patient sample (Fig S1). The differentially expressed genes were strongly associated with the immune response, inflammatory response and innate immune response according to Gene ontology (GO) analysis (Fig 1A). We then mapped the detected lncRNAs to the human genome (UCSC version hg19) and the NONCODE V3.0 database and found an average of 240 up-regulated (range from 164 to 357) and 193 down-regulated (range from 158 to 257) novel lncRNAs in acute-stage influenza patients compared with their recovery-stage counterparts (FC>2, *p<0.05*) and subjected them to a cluster analysis (Fig 1B). To investigate the association between lncRNAs and IAV infection, *in silico* analysis identified 26 novel lncRNAs candidates with notably aberrant expression (Fig 1C). To confirm the expression of these lncRNAs, we separated the neutrophils and monocytes from healthy volunteers and infected the cells with influenza virus A/Beijing/501/2009 (abbreviated as IAV-BJ501 or BJ501) for 12 h and found that XLOC_040025 (*AVAN*) was most significantly up-regulated after IAV-BJ501 challenge (Fig 1D, E). We also infected human alveolar epithelial cells (A549) with IAV-BJ501 and measured the expression of the 26 lncRNAs and found that *AVAN* was most strongly up-regulated after IAV-BJ501 infection (Fig 1F).

**Fig 1.**
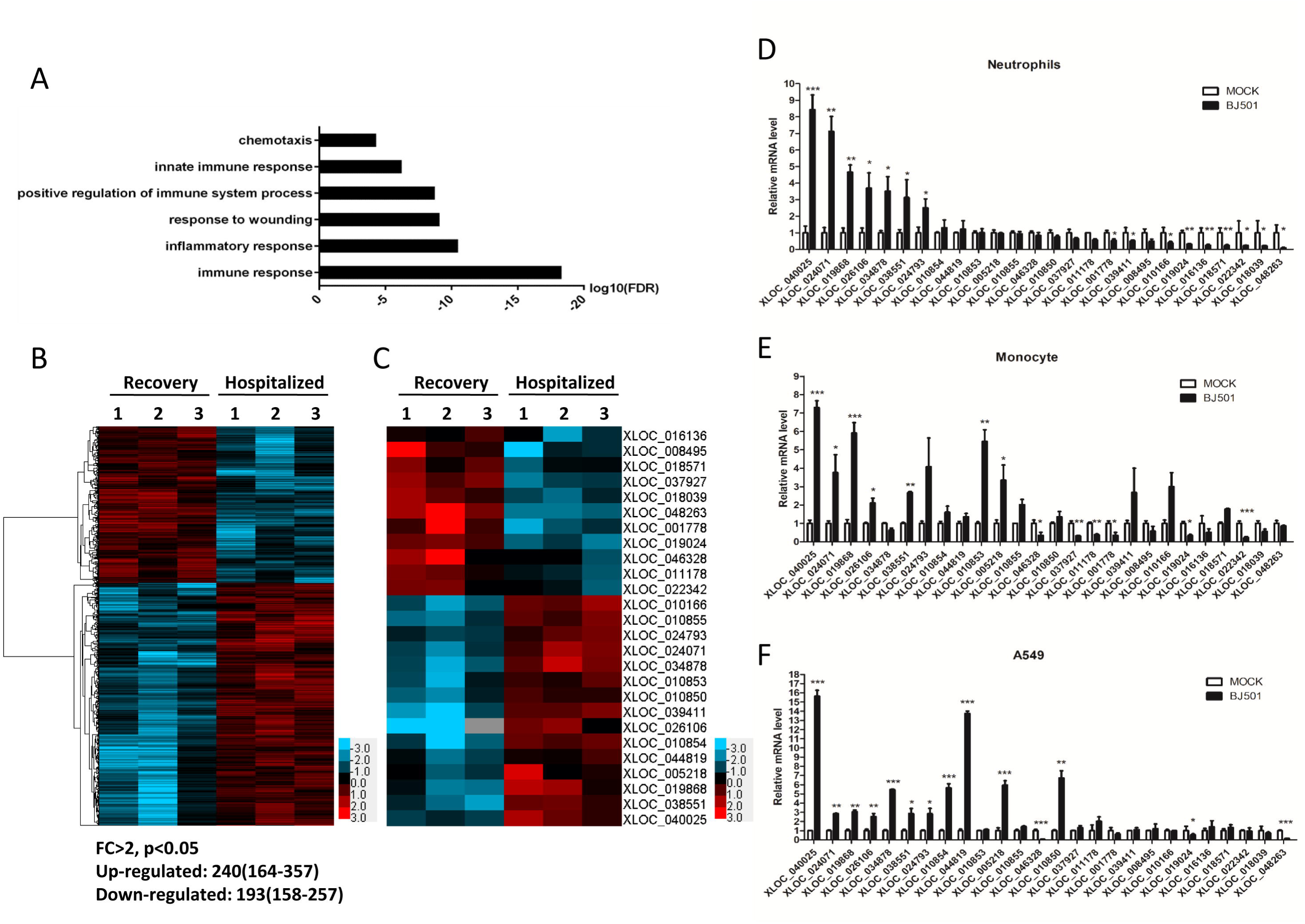
LncRNA expression is regulated by the influenza A virus. (A) Gene ontology (GO) analysis of differentially expressed genes in IAV-infected patient neutrophils compared with their recovery-stage counterparts from RNA-seq data (FC>2; p<0.05). The top six most significantly enriched GO terms are shown. (B) Cluster heat map showing differentially expressed lncRNAs in IAV-infected patient neutrophils compared with recovery-stage samples based on RNA-seq data (FC>2; p <0.05). (C) Cluster heat map showing 26 lncRNA candidates selected via *in silico* analysis from the RNA-seq data (FC>2; p <0.05). (D-E) The expression of 26 lncRNA candidates in neutrophils (D) and monocyte (E) from healthy volunteers stimulated with BJ501 (MOI=0.5) or mock for 12 h by qRT-PCR analysis (n=3; means ± SEM; *p<0.05;**p<0.01;***p<0.001). (F) The expression of 26 lncRNA candidates in A549 cells stimulated with BJ501 (MOI=1) or mock for 24 h by qRT-PCR analysis (n=3; means ± SEM; *p<0.05; **p<0.01; ***p<0.001).

### *AVAN* is preferentially up-regulated following virus infection

We then performed qRT-PCR to quantify *AVAN* expression in the RNA-Seq samples and found that the results were consistent with the RNA-Seq data (Fig S2A). Thus, *AVAN* was chosen for subsequent investigation. To further confirm the expression of *AVAN* in IAV-infected patients, we collected blood samples from 63 additional IAV-positive patients and measured the expression of *AVAN*, which was significantly up-regulated in the neutrophils of these patients (Fig 2A; Table S1). Similarly, *AVAN* was up-regulated in A549 cells after infection with multiple sub-strains of IAV (Fig 2B) and several other RNA viruses, including Sendai virus (SeV) and respiratory syncytial virus (RSV), but not the DNA virus, adenovirus (ADV) (Fig S2B). In addition, Poly (I:C), IFN-β also could significantly stimulate the expression of *AVAN* (Fig S2B). Moreover, *AVAN* expression was significantly up-regulated upon IAV-BJ501 challenge in a time- and dose-dependent manner in A549 cells (Fig 2C, D). Absolute copy number measurement showed that AVAN was expressed at relatively low levels without infection, with 11 or 14 transcript copies in per neutrophil or A549 cell respectively (Figure S2). Additionally, *AVAN* expression was also significantly up-regulated in a time- and dose-dependent manner in the human promyelocytic leukemia cell line (HL60), which is often a substitute for neutrophils, upon IAV-BJ501 challenge (Fig S2B, C). Subcellular fractionation followed by qRT-PCR in IAV-BJ501-infected A549 cells revealed that the amount of *AVAN* localized in the cytoplasm was nearly as same as the one in nucleus (Fig 2E, Fig S2D). Besides, we performed RNA fluorescence *in situ* hybridization (RNA-FISH) with two specific probes in IAV-BJ501-uninfected and –infected A549 cells and found that *AVAN* localized in the cytoplasm and nucleus were almost the same abundance (Fig 2F). In addition, we constructed competitive association FISH to display the specific distribution of *AVAN*. When adding free labeling-probes, the fluorescent dye from biotin labeling-probes was weaken (Fig S2E). *AVAN* exhibits no protein-coding potential according to ORF finder (33) and the coding potential calculator (34) (Fig S2F). Only one transcript variant (approximately 500 nt) of *AVAN* was found in A549 cells, and it was found to be up-regulated upon IAV-BJ501 infection using northern blot analysis (Fig 2G). The exact transcript length (517 nt) of *AVAN* was confirmed by 5’ and 3’ RACE (Table S2), which revealed that *AVAN* is polyadenylated (Fig S2G). Taken together, lncRNA expression profiling of patient neutrophils during IAV infection identified a novel *AVAN* that was preferentially up-regulated in IAV-BJ501-infected cell lines and patient neutrophils upon virus infection.

**Fig 2.**
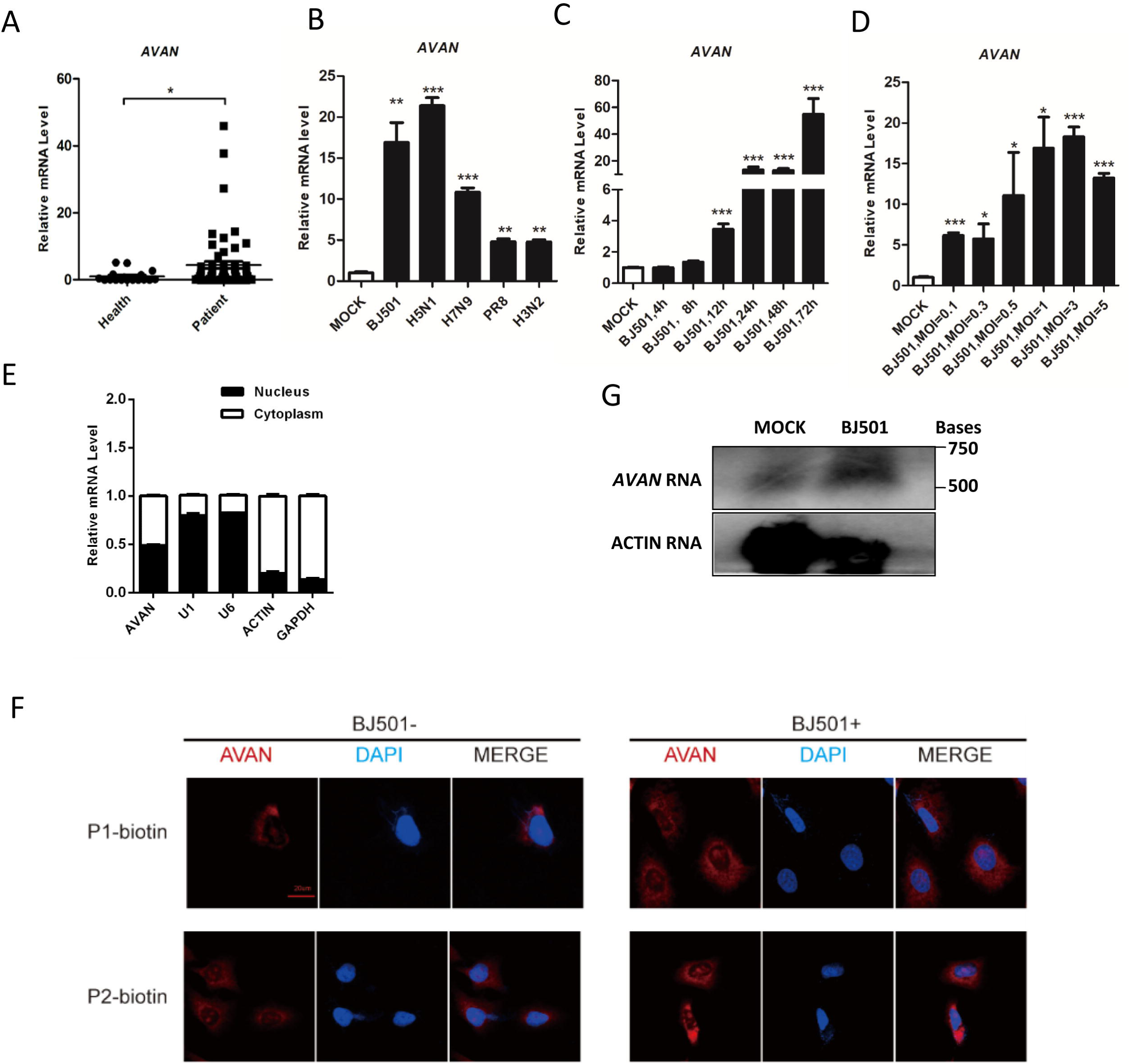
*AVAN* is highly expressed in viral infection. (A) *AVAN* expression in IAV-infected patient neutrophils by qRT-PCR analysis (healthy controls=18; patients=63; *p<0.05). (B) *AVAN* expression in A549 cells infected with BJ501 (MOI=1), H5N1 (MOI=1), H7N9 (MOI=1), PR8 (MOI=1), H3N2 (MOI=1) for 24 h by qRT-PCR analysis (n=3; means ± SEM; **p<0.01; ***p<0.001). (C and D) *AVAN* expression in A549 cells infected with BJ501 at an MOI of 1 for the indicated times (C) or at the indicated MOIs for 24 h (D) by qRT-PCR analysis (n=3; means ± SEM; *p<0.05; **p<0.01; ***p<0.001). (E) Fractionation of BJ501-infected A549 cells followed by qRT-PCR analysis. The U1 and U6 RNAs served as positive controls for nuclear gene expression. The ACTIN and GAPDH RNAs served as positive controls for cytoplasmic gene expression. N, nuclear fraction; C, cytoplasmic fraction. (F) AVAN intracellular localization visualized by RNA-FISH in A549 cells stimulated with MOCK (left) or BJ501 (MOI=1) (right) for 24 h. DAPI, 4’,6-diamidino-2-phenylindole. Probe 1 and 2, AVAN. Scale bar, 10 μm. (G) Northern blotting of *AVAN* in A549 cells treated with mock or BJ501 (MOI=1) at 24h post-infection.

### *AVAN* is essential for antiviral immune responses and neutrophil chemotaxis during IAV-BJ501 infection

To evaluate the potential function of *AVAN*, we next analyzed the related gene coexpression networks from the sequencing data and performed gene set enrichment analysis (GSEA). Influenza viral RNA transcription and replication were negatively related to *AVAN* (Fig S3, Table S3-S8), suggesting that *AVAN* participates in innate antiviral immunity. To identify the functions of *AVAN*, we generated *AVAN*-overexpressing A549 cells (Fig 3A) and found that viral replication was strongly inhibited (Fig 3B). In contrast, virus titer was up-regulated following *AVAN* knockdown with two specific siRNAs (Fig 3H, I), indicating that *AVAN* enhances the regulation of antiviral immune responses.

**Fig 3.**
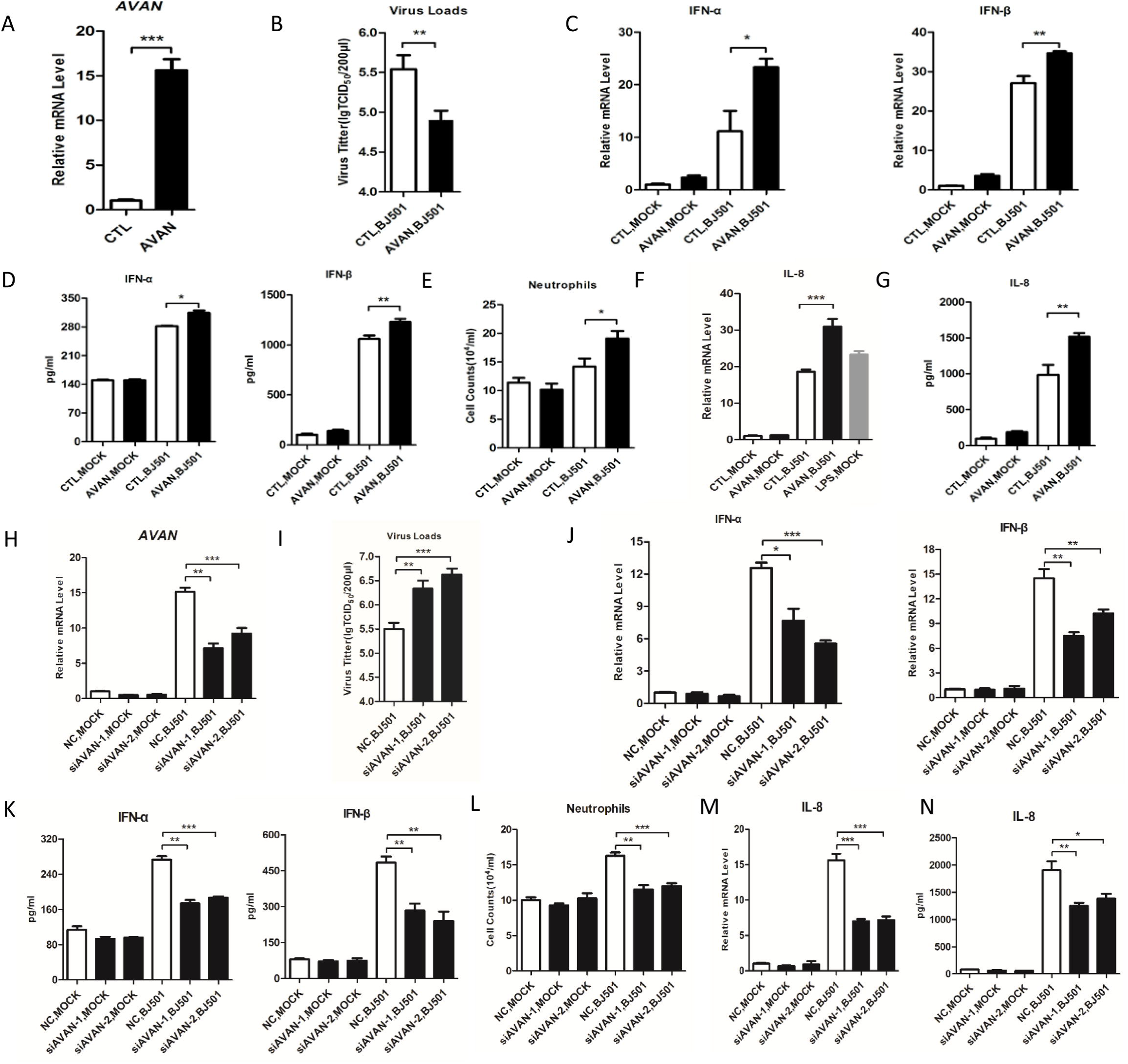
Altered *AVAN* expression has profound effects on type I interferon and neutrophil chemotaxis. (A) The efficiency of *AVAN* overexpression was determined by qRT-PCR in BJ501-infected A549 cells (n=3; means ± SEM; ***p<0.001). (B and I) BJ501 replication in *AVAN*-overexpressing (B) and *AVAN*-knockdown (I) A549 cells examined by the TCID_50_ assay (MOI=1). The virus titers in supernatants were measured at 24 h post-infection (n=3; means ± SEM; **p<0.01; ***p<0.001). (C, D, J and K) IFN-α and IFN-β expression in A549 cells measured by qRT-PCR and ELISA (MOI=1) at 24 h post-infection (n=3; means ± SEM; *p<0.05; **p<0.01; ***p<0.001). (E and L) Transwell assay of neutrophil migration in *AVAN*-overexpressing or *AVAN*-knockdown A549 culture supernatants (n=3; means ± SEM; *p<0.05; **p<0.01; ***p<0.001). (F and M) IL-8 expression in A549 cells were measured by qRT-PCR (MOI=1) at 24 h post-infection (n=3; means ± SEM; ***p<0.001). (G and N) IL-8 expression in A549 cells were measured by ELISA (MOI=1) at 24 h post-infection (n=3; means ± SEM; *p<0.05; **p<0.01). (H) The efficiency of AVAN knockdown was determined by qRT-PCR in BJ501-uninfected or -infected A549 cells (n=3; means ± SEM; **p<0.01; ***p<0.001).

Innate immunity provides the first line of defense against invading pathogens. The recognition of invading virus by pathogen-recognition receptors activates the innate immune system, resulting in the production of type I IFNs (IFN-α and IFN-β) and eliciting antiviral responses (35–37). Thus, we assessed whether *AVAN* regulates virus-induced type I IFN production. The results of qRT-PCR indicate that *AVAN* overexpression increased IFN-α and IFN-β transcript levels upon IAV-BJ501 infection compared with those of the control vector (Fig 3C). To confirm this result, we performed ELISA to measure IFN-α and IFN-β protein expression and found that IFN-α and IFN-β were strongly up-regulated in *AVAN*-overexpressing A549 cells during IAV-BJ501 infection (Fig 3D). Conversely, these phenomena were abolished by knocking down *AVAN* (Fig 3J, K).

The above GSEA data and GO analysis showed that the genes related to *AVAN* were also significantly enriched in chemotaxis and immune cell activation (Fig S4A, Table S9, S10), the main components of innate immune responses. To explore the role of *AVAN* in chemotaxis, we collected culture supernatants of *AVAN*-overexpressing A549cells and performed neutrophil transwell assays, which revealed that *AVAN* overexpression significantly up-regulated neutrophil chemotaxis during IAV-BJ501 infection (Fig 3E). These phenomena were disrupted by knocking down *AVAN* (Fig 3L). The chemokines interleukin-8 (CXCL8, IL-8) is potent neutrophil chemoattractant (38) and is released by a variety of lung cells, including macrophages and epithelial cells as well as neutrophils themselves (39). IL-8 primarily attracts neutrophils and induces them to release lysosomal enzymes, triggering the respiratory burst and increasing the expression of adhesion molecules on the cell surface (40). IL-8 is the primary cytokine involved in the recruitment of neutrophils to the site of damage or infection and plays a crucial role in acute inflammation by recruiting and mediating neutrophils and other cells (41). Thus, we determine whether *AVAN* altered IL-8 expression in A549 cells. *AVAN* overexpression increased IL-8 transcript and protein levels upon IAV-BJ501 infection compared with those of the control vector (Fig 3F, G); this was abolished upon *AVAN* knockdown (Fig 3M, N, Fig S4G, H). To verify our hypothesis, we next overexpressed or knocked down *AVAN* in HL60 cells (Fig S4C, E) and infected them with IAV-BJ501. The cell samples were harvested at the indicated times, and transcript and protein levels of IL-8 were measured. The results from HL60 cells were consistent with those obtained in A549 cells (Fig S4D, F). Meanwhile, we isolated neutrophils from human primary cell and found that *AVAN* could stimulate the IL-8 transcript upon IAV-BJ501 infection (Fig S4I, J), and this was abolished upon *AVAN* knockdown (Fig S4K, L). Collectively, these data indicate that lncRNA-*AVAN* plays a vital role in antiviral responses by positively regulating type I IFN induction and neutrophil chemotaxis.

### *AVAN* up-regulates ISG and chemokine expression

To verify the global influence of *AVAN* during virus infection and to gain further insight into *AVAN* activity, we performed cellular transcriptome profiling on A549 cells using cDNA microarrays and found that *AVAN* overexpression altered the expression of 81 genes after 14 h of BJ501 infection compared with control cells (Fig 4A, Table S11). Most of these divergent genes were associated with antiviral innate immune responses and inflammatory diseases according to Reactome pathway analysis (Fig 4B) and FunDO diseases analysis (Fig S5). Notably, genes associated with immune system IFN signaling and cytokine signaling were markedly up-regulated in cells with ectopic *AVAN* expression compared with control cells, including chemokines and ISGs (IFN stimulated genes) (Fig 4C). These significantly up-regulated genes, including MX1, ISG15, IFIT2, OASL, IFIM3, TNFAIP3, CXCL2 and CCL5, were confirmed by qRT-PCR (Fig 4D), and these were abolished after *AVAN* knockdown upon IAV-BJ501 infection (Fig 4E). These data again reveal that *AVAN* strongly participates in antiviral innate immune processes and neutrophil chemotaxis.

**Fig 4.**
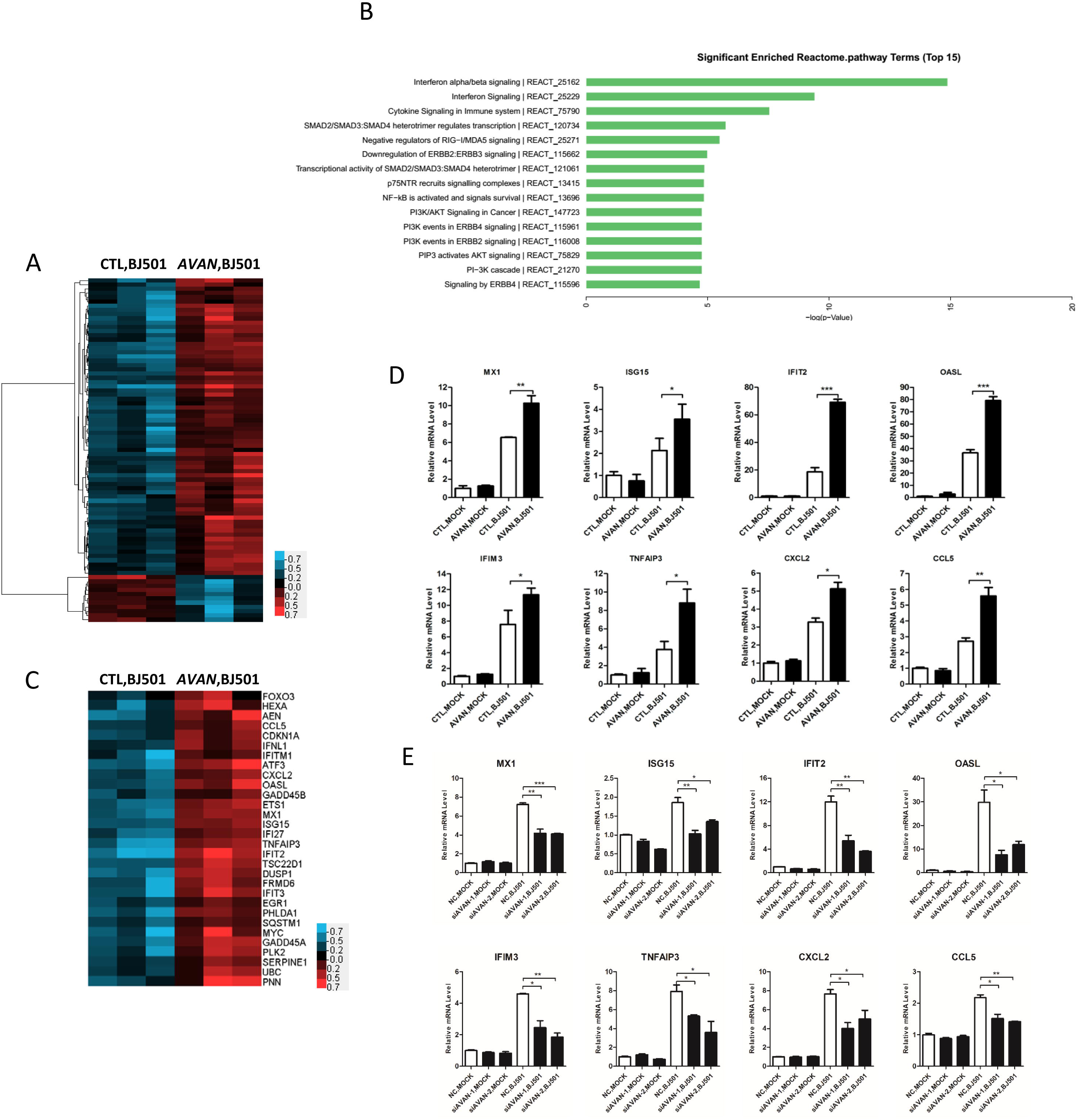
*AVAN* up-regulates ISG and chemokine expression. (A) Cluster heat map showing altered mRNA expression in *AVAN*-overexpressing A549 cells and EV control A549 cells infected with BJ501 (MOI=1) for 14 h by cDNA microarray (n=3; FC>1.3; p<0.05). (B) Reactome pathway analysis of the altered mRNA expression from the cDNA microarray analysis. (C) Cluster heat map showing the altered expression of mRNAs involved with ISGs and cytokine mRNAs that were up-regulated during *AVAN* overexpression. (D and E) The mRNA levels of selected genes in *AVAN*-overexpressing or -konckdown A549 cells and control A549 cells infected with BJ501 (MOI=1) for 14 h by qRT-PCR analysis (n=3; means ± SEM; *p<0.05; **p<0.01; ***p<0.001).

### *AVAN* triggers FOXO3a expression in the nucleus to enhance neutrophil chemotaxis

Recent studies have shown that divergent lncRNAs can positively regulate the transcription of nearby genes through chromatin remodeling (4,42–46). The genomic location of *AVAN* suggested that it was divergently transcribed from a position 500-bp upstream of the FOXO3a gene (Fig S2G). Interestingly, mRNA array data revealed that FOXO3a is significantly up-regulated following *AVAN* overexpression during IAV-BJ501 infection (Fig 4C). To investigate the role of *AVAN* in FOXO3a expression, we measured the abundances of FOXO3a after overexpressing or knocking down *AVAN* in A549 and HL60 cell lines. The overexpression of *AVAN* up-regulated FOXO3a transcription in A549 cells, consistent with its role in enhancing FOXO3a protein expression (Fig 5A; Fig S6A left). Conversely, knocking down *AVAN* down-regulated FOXO3a transcription and protein expression (Fig 5B; Fig S6A right). In addition, the results in HL60 were consistent with these observed in A549 cells (Fig S6B).

**Fig 5.**
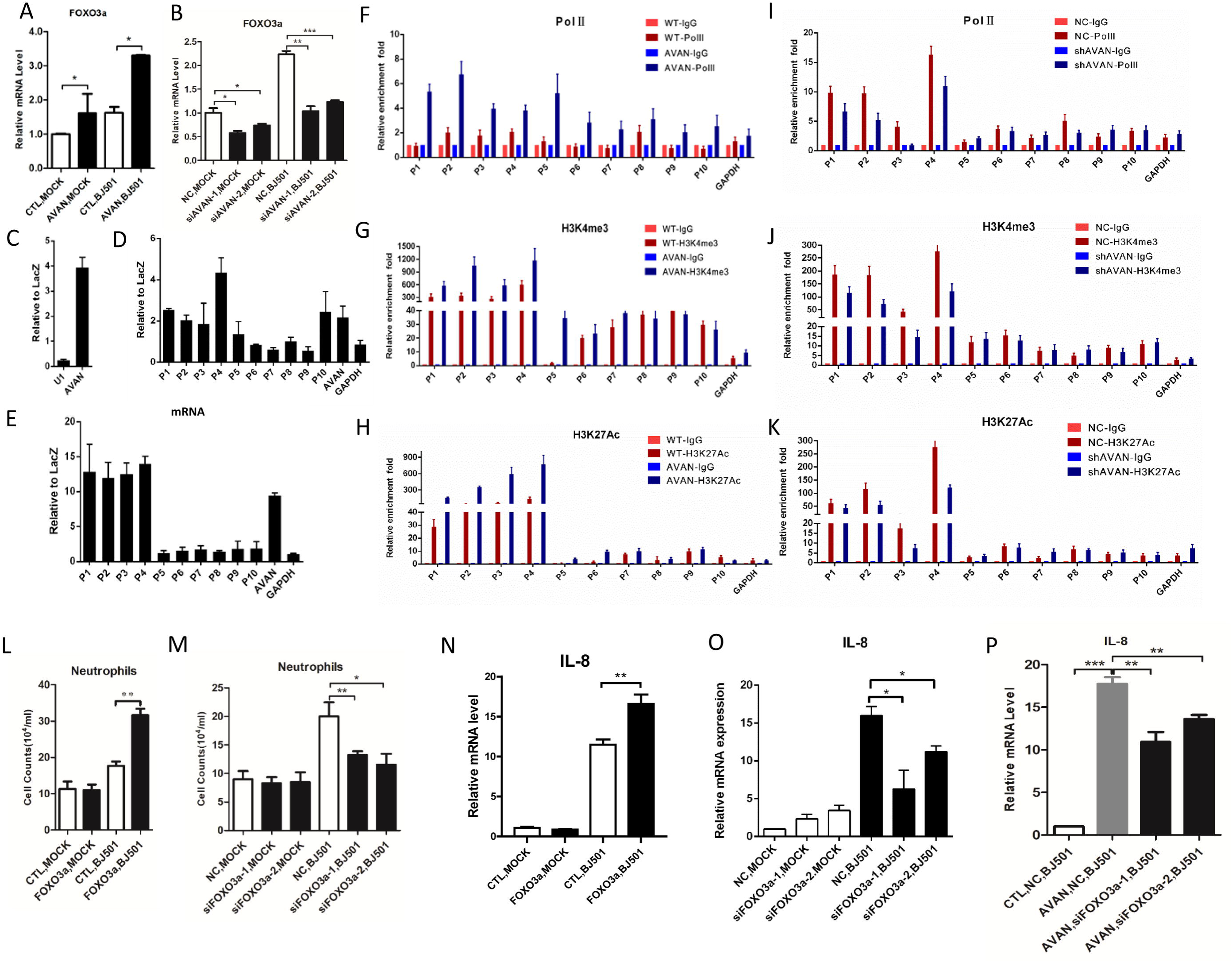
*AVAN* enhances FOXO3a expression *in nucleus*. (A and B) FOXO3a expression during virus infection in A549 cells were measured by qRT-PCR. A549 cells were transfected with *AVAN* plasmids (A) or siRNA (B) (n=3; means ± SEM; *p<0.05; **p<0.01; ***p<0.001). (C) Enrichement of AVAN in ChIRP assay analyzed by qRT-PCR, U1 as a negative control. (D and E) ChIRP assay showing that AVAN binds directly to the FOXO3a promoter DNA in BJ501-uninfected (D) and -infected (E) A549 cells. P1-P10 represent the different regions in FOXO3a promoter or gene body locus. (F-K) H3K4me3 and H3K27ac levels and Pol II binding of the FOXO3a promoter were analyzed through ChIP followed by qRT-PCR in BJ501-infected A549 cells. (L and M) Transwell assay of neutrophil migration in FOXO3a-overexpressing or FOXO3a-knockdown A549 culture supernatant (n=3; means ± SEM; *p<0.05; **p<0.01). (N and O) IL-8 expression in FOXO3a-overexpressing (N) or FOXO3a-knockdown (O)A549 cells were measured by qRT-PCR (n=3; means ± SEM; *p<0.05; **p<0.01). (P) IL-8 expression in *AVAN*-overexpressing or/and FOXO3a-knockdown A549 cells were measured by qRT-PCR (n=3; means ± SEM; **p<0.01; ***p<0.001).

The association of *AVAN* with FOXO3a expression and the subcellular localization of *AVAN* in both the cytoplasm and nucleus (Fig 2E, F) prompted us to test whether *AVAN* interacts with the FOXO3a promoter. To this end, we conducted chromatin isolation by RNA purification (ChIRP) using antisense oligoes against *AVAN* followed by qPCR in BJ501 -uninfected and -infected A549 cells and found that *AVAN* bound to sequences upstream of FOXO3a (Fig 5C, D, E; Fig S6C). We further verified the potential role of *AVAN* in modulating chromatin modifications at the FOXO3a promoter region. Chromatin immunoprecipitation (ChIP)-qPCR revealed that ectopic *AVAN* expression increased Pol II binding, as well as H3K4me3 and H3K27ac levels, at the *AVAN*-coated FOXO3a promoter in virus infected A549 cell (Fig 5F, G, H) and uninfected A549 cell (Fig S6D, E, F). In contrast, knocking down *AVAN* decreased Pol II binding and H3K4me3 and H3K27ac accumulation (Fig 5I, J, K; Fig S6G, H, I).

Chemokines were markedly up-regulated in cells with ectopic *AVAN* expression (Fig 4C). A previous study reported that FOXO3a can regulate chemokine expression (47). To further investigate whether *AVAN*-promoted chemokines expression is associated with FOXO3a, we measured IL-8 expression in FOXO3a overexpressing A549 cells. Ectopic FOXO3a expression significantly promoted IL-8 expression and neutrophil chemotaxis during IAV-BJ501 infection (Fig 5L, N; Fig S6J). In contrast, knocking down FOXO3a decreased IL-8 expression and neutrophil chemotaxis during IAV-BJ501 infection (Fig 5M, O; Fig S6K). In addition, we knocked down FOXO3a in *AVAN*-overexpressing A549 cells and found that knockdown abrogated *AVAN*-induced effects on IL-8 expression during IAV-BJ501 infection (Fig 5P). These data provide evidence that *AVAN* plays a critical role in regulating FOXO3a expression in nucleus by associating with the FOXO3a promoter and performing chromatin remodeling, which additionally increased IL-8 and neutrophil chemotaxis.

### *AVAN* direct binds to TRIM25 and enhances the TRIM25-mediated activation of RIG-I signaling

Previous experiments demonstrated that *AVAN* is located in the cytoplasm (Fig 2E, 2F). To investigate the molecular mechanism of *AVAN* in the cytoplasm, RNA pull-down assays using biotin-labeled *AVAN* or *AVAN* antisense control followed by mass spectrometry (MS) analysis were performed. The result of MS revealed that E3 ubiquitin ligase TRIM25, an RNA-binding protein (48, 49), was pulled down by *AVAN* in IAV-BJ501-infected A549 cells but not by the *AVAN* antisense control (Fig 6A, Fig S7A). This result was confirmed by *AVAN* RNA pull-down western blot experiments (Fig 6B). Besides, ChIRP followed by western blot revealed that *AVAN*-specific probes could pull down TRIM25 while LacZ couldn’t (Fig 6C). To validate the interaction between *AVAN* and TRIM25, we immunoprecipitated TRIM25 from IAV-BJ501-infected A549 cells and quantified the protein-bound *AVAN*. Significantly higher levels of *AVAN* were detected with exogenous (Fig 6D) and endogenous (Fig 6E) TRIM25 immunoprecipitation than with the isotype immunoglobulin G (IgG) control. Furthermore, we constructed Flag-tagged TRIM25 truncated proteins containing the SPRY domain, B box/central coiled-coil domain (CCD), or RING domain and performed *AVAN* RNA pull-down western blot experiments. The results showed that only the B box/central CCD domain was involved in the *AVAN*-TRIM25 association (Fig 6F). Furthermore, to explore the TRIM25 binding site on AVAN RNA, three truncated probes from AVAN were used for RNA pull-down assay. The result indicated that the TRIM25-binding activity mapped between nucleotides 1 and 200 (Fig 6G). Together, these findings indicate that *AVAN* physically interacts with TRIM25 and that the B box/central CCD of TRIM25 and 1-200nt of AVA*N* contribute to this association.

**Fig 6.**
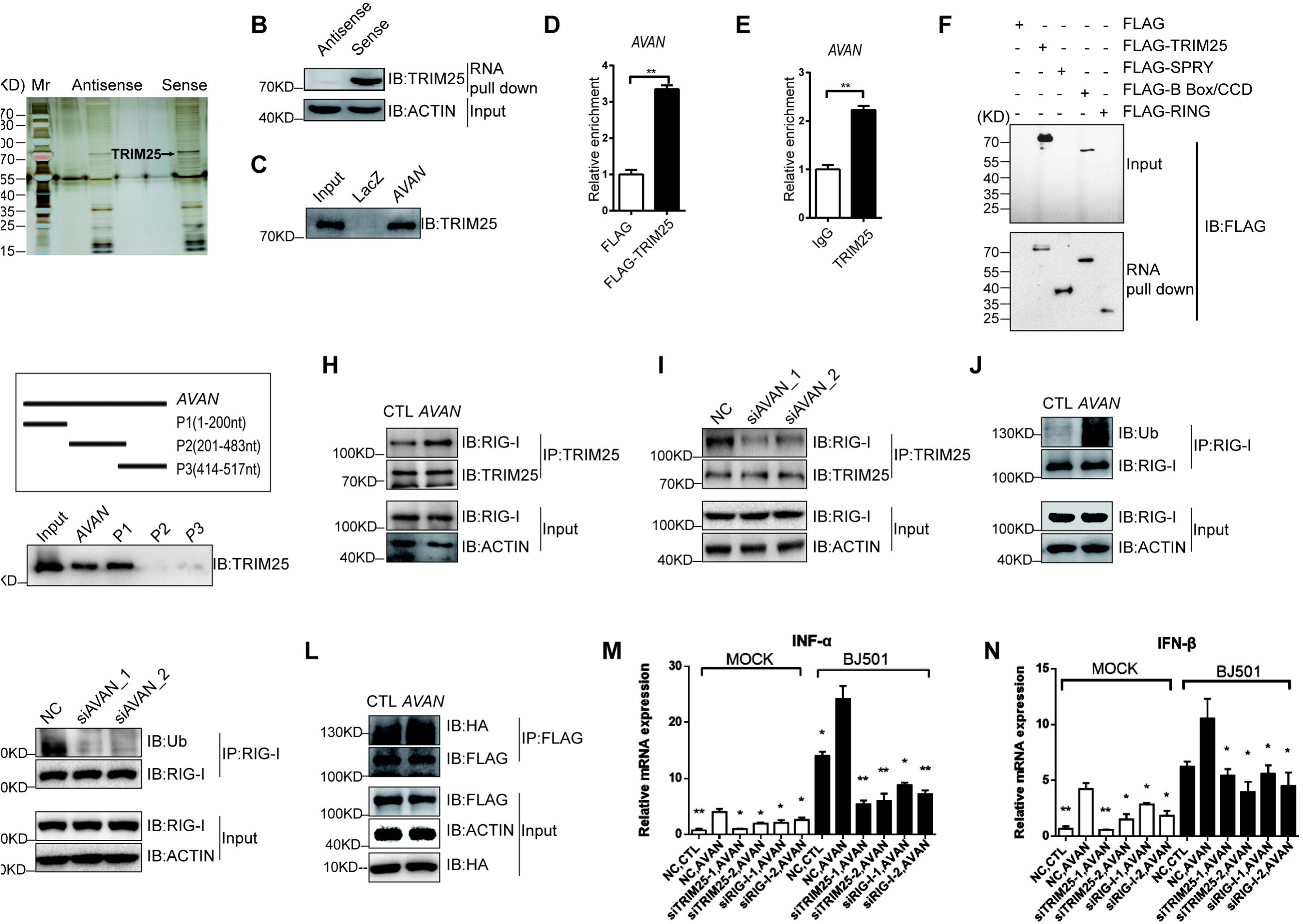
*AVAN* direct binds to TRIM25 and enhances the antivirus immune response. (A) RNA pull-down of *AVAN*-associated proteins using biotinylated AVAN or antisense probes. Isolated proteins were resolved by SDS-PAGE followed by silver staining. (B) Pull-down western blot showing that *AVAN* can bind directly to TRIM25. (C) ChIRP followed by western blot show that *AVAN* can bind to TRIM25. (D and E) Exogenous (D) and endogenous (E) RIP of TRIM25 in BJ501 infected cells using anti-TRIM25 or anti-IgG antibodies. The relative enrichment fold of *AVAN* was calculated by qRT-PCR. (F) *AVAN* pull-down western blot with lysates of A549 cells transfected with Flag, Flag-TRIM25, Flag-SPRY, Flag-B Box/CCD or Flag-Ring. (G) Truncated *AVAN* pull-down, truncates (upper panel) were obtained via *in vitro* transcription and incubated with BJ501-infected A549 lysates for RNA pulldown. (H and I) TRIM25 co-immunoprecipitation with proteins from lysates of BJ501-infected A549 cells transfected with *AVAN*s or siRNAs, followed by immunoblotting. Anti-TRIM25 and anti-RIG-I antibodies were used for immunoprecipitated. (J and K) Immunoblot analysis of endogenous RIG-I ubiquitylation in BJ501-infected A549 cells transfected with *AVAN*s or siRNAs. Anti-RIG-I antibody was used for immunoprecipitated. (L) Immunoblot analysis of proteins immunoprecipitated with anti-Flag from lysates of BJ501-infected A549 cells transfected with *AVAN*, HA-Ub and Flag-tagged RIG-I. (M and N) IFN-alpha (M) and IFN-beta (N) expression upon AVAN transfection in A549 cells that were infected by BJ501 or not (MOI=1) at 24h post-infection, and then individually knock down RIG-I or TRIM25, analyzed by qRT-PCR (n=3; means ± SEM; *p<0.05; **p<0.01).

To test whether *AVAN* enhances the IFN-mediated antiviral innate immune response through TRIM25 and RIG-I signaling, we knocked down endogenous TRIM25 in HEK-293T cells and observed a markedly reduced effect of ectopic *AVAN* expression on the activation of IFNB1-responsive reporters in the context of IAV-BJ501 infection (Fig S7B). RIG-I is polyubiquitinated by TRIM25, which attaches K63-linked polyubiquitin to the sensor (50), Previous study showed that a stabilization of TRIM25-RIG-I interaction is important for a sustained antiviral IFN response (51). We next examined the effect of *AVAN* on the interaction between TRIM25 and RIG-I. We found that ectopically expressing *AVAN* markedly enhanced the association between TRIM25 and RIG-I upon IAV-BJ501 infection, and knockdown of AVAN weakened this association (Fig 6H, I). Additionally, different fragments of *AVAN* were overexpressed in A549. We observed that P1 (1-200nt) could enhance the association between TRIM25 and RIG-I, and significantly upregulate the expression of IFN-α and IFN-β, as full-length *AVAN*, while fragment P2+P3 (201-517nt) couldn’t (Fig S7C, D). Moreover, endogenous or exogenous RIG-I ubiquitylation was significantly increased in *AVAN*-overexpressing cells in the context of IAV-BJ501 infection compared with the control vector while RIG-I ubiquitylation was decreased in AVAN-knockdown cells (Fig 6J, K, L). Furthermore, we explored the impact of altered *AVAN* levels on RIG-I signaling. Total RIG-I protein levels significantly upregulated between *AVAN*-overexpressing and control cells. In addition, *AVAN* overexpression increased TBK1 and IRF3 phosphorylation upon IAV-BJ501 infection (Fig S7E, left). In contrast, *AVAN* knockdown abolished these changes (Fig S7E, right). To test whether AVAN enhances the IFN-mediated antiviral innate immune response through TRIM25 and RIG-I signaling, we performed qPCR analysis measuring IFN-alpha/beta upon AVAN transfection, and then individually knock down RIG-I or TRIM25. The results showed that knock down RIG-I or TRIM25 abrogated AVAN-induced effects on IFN-alpha/beta expression during IAV-BJ501 infection (Fig 6M, N, Fig S7F, G). Together, these data show that *AVAN* can enhance the interaction between TRIM25 and RIG-I and the ubiquitylation of RIG-I in the cytoplasm. Thus, we conclude that *AVAN* promotes TRIM25- and RIG-I-mediated antiviral innate immune signaling.

### AVAN expression significantly alleviates IAV-BJ501 virulence in transgenic mice

Although we fail to detect mouse lncRNA, mouse genome contains the *AVAN* homolog sequences. To further investigate the *in vivo* effect of *AVAN* on IAV-BJ501 infection, transgenic mice (TG) mice expressing human *AVAN* were generated as previously described(52). TG mice lung with high *AVAN* expression were identified and selected for this experiment (Fig 7A). TG mice and WT littermates were i.n. inoculated with BJ501 virus. As expected, IAV-BJ501 displayed a considerably lower virulence in TG mice than that in WT mice. The survival rates were 60% and 20% in TG mice and WT mice after 14 days post-infection, respectively (Fig 7B). The TG mice exhibited significantly reduced body weight loss at 7 days post-infection compared with the WT mice (Fig 7C). Meanwhile, we compared the lung edema and viral loads of infected TG mice with WT littermates and found that lung wet-to-dry ratio and viral titer in TG mice was significantly lower than that in WT mice (Fig 7D, E). Pathologic examination by HE attaining revealed that histopathlogic alterations improved in TG mice (Fig 7F). Subsequently, we detected FOXO3a expression levels and found that FOXO3a expression were undetectable in TG mice (data not shown), likely due to low *AVAN* homology in humans and mice. Striking, we found that *AVAN* also bind to rodent TRIM25 through RNA pull-down using TG mice tissue (Fig 7G) and promote type I IFN expression *in vivo* (Fig 7H). Similarly, we constructed an *AVAN*-containing AAV2/9 vector and a control vector, which were then delivered into 4-week-old C57L/B6 mice via intranasal (i.n.) administration. We found that *AVAN* was ectopically expressed in the lung of AAV2/9-*AVAN* treated mice (Fig S8A). Strikingly, after IAV-BJ501 infection, the groups pretreated with AAV2/9-*AVAN* exhibited significantly increased survival rates and reduced body weight loss at 10 days post-infection compared with the control group (Fig S8B, C). Furthermore, lung edema, measured as the wet-to-dry ratio of whole lung, was ameliorated (Fig S8D), and improved lung histopathology was observed in infected mice pretreated with AAV2/9-*AVAN* (Fig S8E, F). Moreover, the virus titer from the lungs of infected mice was also significantly reduced in AAV2/9-*AVAN*-treated mice compared with that in control mice (Fig S8G, H). In addition, AAV2/9-*AVAN* promote type I IFN expression *in vivo* (Fig S8I). Taken together, these observations indicate that expression of lncRNA *AVAN* can ameliorate IAV-BJ501-induced acute lung injury *in vivo* and protect mice from IAV-BJ501 infection.

**Fig 7.**
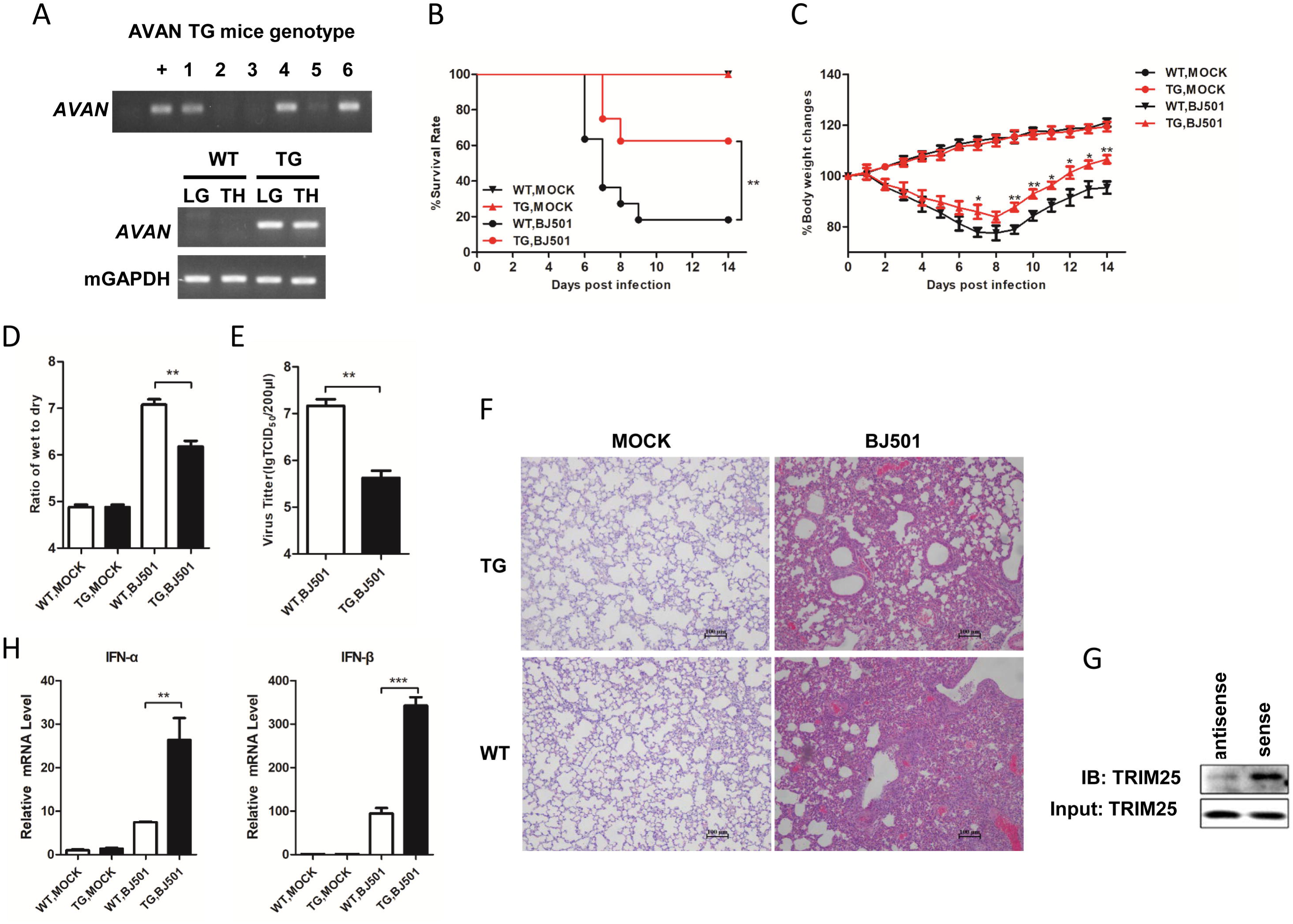
AVAN expression significantly alleviates IAV virulence in transgenic mice. (A) The genotype of C57BL/6J TG mice were identified by PCR of mouse tail DNA and RT-PCR of tissue RNA. LG: lung; TH: Thymus. (B and C) Four-week-old TG and WT mice were inoculated with 10^5.125^ TCID_50_ BJ501 virus, respectively. Survival rates and body weight changes of wild-type mice (n=10 for each group) were monitored for 2 weeks following BJ501 challenge. (D) Wet to dry ratio of lung tissues (n=5 for each group) at 5 DPI. (E) Viral load of lung tissues (n=5 for each group) at 5 DPI. (F) HE-stained images and infiltrating cell counts (n=100 fields) in lung tissues at 5 DPI (magnification=200×). (G) Immunoblot analysis showed that *AVAN* can bind to TRIM25 *in vivo*. (H) IFN-α and IFN-β expression in the lung of TG mice measured by qRT-PCR. *p<0.05, **p<0.01 and ***p<0.001.

## Discussion

Recently, increasing evidence supports the importance of lncRNAs in host-virus interactions, and lncRNAs are an emerging paradigm in the regulation of innate immune responses. Previous studies have described several lncRNAs that are differentially expressed during viral infection, including in severe acute respiratory syndrome coronavirus (SARS-CoV)-infected mice, IAV-infected human lung cells and enterovirus 71 (EV71)-infected rhabdomyosarcoma (RD) cells (53–55). However, little is known about the lncRNA profile of influenza patients. In this study, we profiled the transcriptome of neutrophils isolated from IAV-infected patients via RNA-Seq technology for the first time. Through a comprehensive analysis of these data, we identified 404 differentially expressed mRNAs and 433 differentially expressed noncoding RNAs in neutrophils of influenza patients. Among these highly expressed genes, we found an lncRNA named *AVAN* that plays a pivotal role in antiviral responses.

Several studies have shown that differentially expressed lncRNAs function as negative or positive regulators in various critical steps of the antiviral response (56). For example, BISPR lncRNA can regulate the antiviral IFN response through the induction of the expression of the genomically neighboring gene BST2 *in cis*. Another IAV-associated intronic antisense lncRNA, negative regulator of antiviral response (NRAV), modulates antiviral responses by suppressing ISG transcription via altered histone modifications (52). A recent study show that lnc-Lsm3b can compete with viral RNAs in the binding of RIG-I monomers and involve in RIG-I-mediated antiviral response (57). In contrast, Herein, our data displayed that *AVAN* acts as a positive regulator in the antivirus response through two different mechanisms. In one hand, cytoplasmic lncRNA-AVAN directly binds TRIM25 and enhances the association of TRIM25 and RIG-I and the ubiquitylation of RIG-I, thereby promoting TRIM25- and RIG-I-mediated antiviral innate immune signaling. In the other hand, nuclear lncRNA-*AVAN* positively regulates the transcription of forkhead box O3A (FOXO3a) *in nucleus* by associating with its promoter and inducing chromatin remodeling to enhance neutrophil chemotaxis.

FOXO3a is a member of the Forkhead box O (FoxO) family of transcription factors, which is a PTEN/PI3K/AKT effector functioning in diverse cellular activities such as the induction of cell-cycle arrest, stress resistance, apoptosis, differentiation, and metabolism (58–60). FOXO3a reportedly functions as a major transcriptional regulator in the maintenance of neutrophil activation during inflammation (61). However, little is known about the function of FOXO3a during the immune response in epithelial cells. Previous research demonstrated that FOXO3a repressed cytokine expression induced by LPS or bacteria (47). In this study, FOXO3a plays a different function during IAV infection. FOXO3a promotes IL-8 expression in IAV-BJ501-infected A549 cells, which additionally enhances neutrophil chemotaxis. *AVAN* is located 1449-bp upstream of FOXO3a. We found that lncRNA-*AVAN* can markedly increase FOXO3a expression through chromatin remodeling in A549 cells. Additionally, *AVAN* can enhance neutrophil chemotaxis by upregulating the expression of IL-8. However, knockdown FOXO3a blocked the upregulation of IL-8 by *AVAN*. In the other hand, to confirm how much of the rise in the level of FOXO3a after AVAN overexpression is the result of enhanced IFN response, as opposed to a direct impact by AVAN on FOXO3a locus, we also used HL60 cells to evade IFN effects (data not shown here). Meanwhile, as seen in Figure 5, our data showed that there is a direct effect of AVAN on FOXO3a promoter and thereby enhanced Pol II binding and H3K4me3 and H3K27ac accumulation. Of course, we would generate a mutant AVAN that can induce IFN response similar to WT AVAN but can’t bind the FOXO3a promoter or change its expression level in subsequently studies. In addition, our results exhibited that neutrophils were isolated from fresh peripheral blood and were infected in vitro with influenza virus does not mimic their actual status in vivo. How and where AVAN is induced in neutrophils need to be further investigated. Thus, we propose that *AVAN* may enhance neutrophil chemotaxis through the regulation of FOXO3a expression. However, the detailed mechanism by which FOXO3a regulates the expression of chemokines requires further exploration.

Retinoic acid-inducible gene-I (RIG-I)-like receptors (RLRs) play a well-known role in RNA virus recognition (62). Previous reports have shown that RIG-I is the main host sensor involved in recognizing cytoplasmic viral RNA and subsequently initiating immune responses to eliminate infection. TRIM25, an E3 ubiquitin ligase, is critically important in the RIG-I signaling pathway. TRIM25 interacts with the CARD1 region of RIG-I and enhances RIG-I polyubiquitylation to initiate RIG-I-mediated antivirus responses (50). A previous study found that viruses have evolved sophisticated mechanisms to evade the host immune response through regulating the role of TRIM25. IAV nonstructural protein 1 (NS1) specifically inhibits TRIM25-mediated RIG-I CARD ubiquitination, thereby suppressing the RIG-I signaling pathway (63). Our results demonstrated that the host can also regulate TRIM25 to increase the immune response. *AVAN* can directly bind with the B box/central CCD of TRIM25 in A549 cells. Furthermore, the interaction of *AVAN* with TRIM25 promotes the association between TRIM25 and RIG-I and enhances RIG-I ubiquitylation, thereby promoting IFN expression. It is worth noting that *AVAN* binds TRIM25 and facilitates IFN expression in mice. In addition, we explored the impact of AVAN overexpression or knockdown in IFN induction, chemo attraction of neutrophils or expression of cytokines and chemokines (Fig 3), speculated that lncRNA AVAN is critical in the IFN induction pathway. Although Cadena et al. reported that TRIM25 is not E3 ligase relevant to RIG-I (64), likely reason is related to various RNA virus subtypes. Of course, additional experiments were need to solidify the TRIM25-RNA interaction. To the best of our knowledge, our work represents the first identification of lncRNA-*AVAN* as a novel partner of TRIM25 and demonstrates that *AVAN* suppresses IAV-BJ501 replication via TRIM25-RIG-I-dependent antiviral pathways.

In conclusion, we provide the first lncRNA landscape of IAV-infected patient neutrophils and identify the function of *AVAN*, a novel lncRNA, in the innate antiviral immune response. *AVAN* can enhance the chemotaxis of neutrophils by promoting FOXO3a expression *in nucleus.* On the one hand, *AVAN* can serve as a positive regulator of RIG-I signaling by directly binding TRIM25 and enhancing the association between TRIM25 and RIG-I *in cytoplasm*. (Fig 8). Of course, the mechanism of action of *AVAN in vivo* (transgenic mice) during IAV-BJ501 and its role in infection by other viruses warrant future investigations. Understanding the powerful roles of ubiquitous and versatile lncRNAs and identifying these IAV-related lncRNAs, especially the novel lncRNA-*AVAN*, may provide insights into the pathogenesis of viral infection, with lncRNA-*AVAN* potentially representing an essential target for intervention.

**Fig 8.**
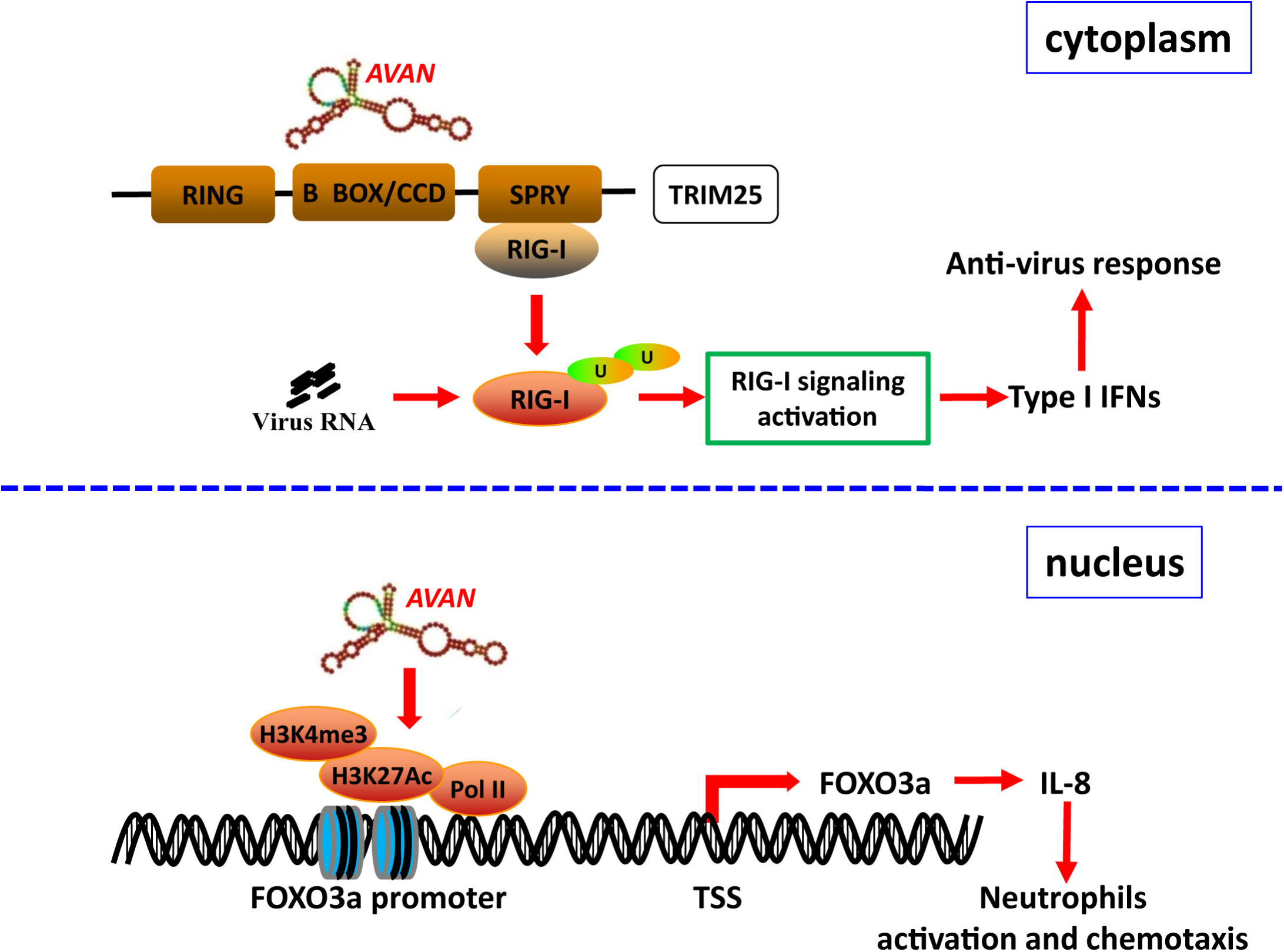
Schematic of the mechanisms by which *AVAN* regulates antivirus responses and neutrophil chemotaxis.

## Materials and methods

### Cells and viruses

A549 cells were cultured in DMEM/F12 (1:1) supplemented with 10% (v/v) FBS and penicillin-streptomycin (100 U/ml); 293T cells were cultured in DMEM/High Glucose supplemented with 10% (v/v) FBS and penicillin-streptomycin (100 U/ml); and HL60 and THP-1 cells were cultured in 1640 supplemented with 10% (v/v) FBS and penicillin-streptomycin (100 U/ml). The influenza virus A/Beijing/501/2009 (abbreviated as IAV-BJ501 in manuscript or BJ501 in Figures) and SeV used in this study were propagated by inoculation into 9 to 11-day-old specific pathogen-free (SPF) embryonated chicken eggs via the allantoic route as described previously. Virus stocks were aliquoted and stored at -80°C until use.

### Mice

Pathogen-free 4-6-week-old female C57BL/6 mice were purchased from the Laboratory Animal Center, AMMS, Beijing, China. All procedures, including animal studies, were conducted following the National Guidelines for the Care of Laboratory Animals (2006-398) and performed in accordance with institutional regulations after protocol review and approval by the Institutional Animal Care and Use Committee of the Academy of Military Medical Sciences (project no. 2012-005). Mice were lightly anesthetized and subjected to i.n. inoculation with a tissue culture infective dose (TCID) of 10^5^ TCID50 of BJ501 influenza virus in a volume of 20 μl per mouse. Control mice were inoculated with 20 μl of allantonic fluid. Inoculated mice were maintained under SPF conditions and monitored daily for weight loss and mortality or infection for 14 days post-infection. Survival rate, body weight changes, histological examination and acute lung edema (wet-to-dry ratio) were determined as described previously (65, 66). The number of infiltrating inflammatory cells was counted and presented as the number of cell per 200× field.

### Transgenic mice and virus challenge

The AVAN transgenic C57BL/6 mice were generated as previously described. TG mice were i.n. inoculated with BJ501 virus. Mouse lungs were collected for viral loads, wet to dry ratio and pathology. All mice experiments were approved by the Ethics Committee of Beijing Institute of Microbiology and Epidemiology.

### RNA interference

A549, HL60 and primary human neutrophil cells were transfected with siRNA targeting *AVAN*, FOXO3a, TRIM25 and RIG-I using jetPRIME (Polyplus) according to the manufacturer’s instructions. Two ASOs were also used to knockdown *AVAN* in A549 cells. The siRNA and ASO sequences used in the experiments are listed in Supplementary Table S12.

### Neutrophils and monocyte isolation

Neutrophils and monocyte were isolated from fresh peripheral blood using Percoll PLUS (GE healthcare) according to the instructions supplied by the manufacturer. For neutrophils collection, briefly, fresh peripheral blood was taken from IAV patients and healthy human donors and layered on a 2-step Percoll PLUS gradient (75% and 60%), then centrifuged at 500×g for 25 minutes. The cells at the interface between 75% and 60% were collected and washed twice with PBS and re-suspended in RPMI-1640 medium (Gibco) supplemented with 10% FBS and 100 U/ml penicillin-streptomycin at 37°C in a humidified atmosphere of 5% (v/v) CO_2_.

### ELISA

Cytokines levels were measured using an ELISA kit (Dakewe, Beijing).

### Neutrophil chemotaxis assay

*In vitro* chemotactic assays were performed in 24-well Millicell hanging-cell culture inserts (Millipore). Briefly, the cells were starved by incubation for 18-24 h prior to the assay in serum-free RPMI 1640 medium and then washed twice with sterile serum-free medium containing 0.5% BSA. The bottom wells were loaded with the supernatant from *AVAN*-overexpressing or knockdown A549 cells 24 h after BJ501 or PBS treatment to a final volume of 300 μl. The top wells were loaded with neutrophils (10^6^cells/ml; 250 μl from RPMI suspension). The top and bottom wells were separated by a porous membrane (3-μm pore size). A cover plate was added, and the cells were incubated for 1 h at 37°C with 5% CO_2_. At the end of the incubation period, the top wells were removed, and the number of cells in the bottom wells was counted using a cell counter. The results are expressed as relative neutrophil migration (number of cells from the tested group/number of cells from the corresponding control vehicles).

### Antibodies and reagents

The primary antibodies anti-RIG-I(D14G6), anti-TBK1(D1B4), anti-phospho-TBK1 (Ser172, D52C2), anti-IRF3(D6I4C), anti-phospho-IRF3 (Ser396,4D4G), anti-β-Actin(13E5), anti-FOXO3a, anti-H3K4me3 and anti-rabbit IgG, were purchased from Cell Signaling Technology. Anti-Flag, anti-HA, anti-H3K27ac and anti-TRIM25 were purchased from Sigma. Anti-Pol II was purchased from Abcam. Streptavidin C1 Beads were purchased from Invitrogen. Protein A/G PLUS-Agarose were purchased from Santa Cruz. Protein A/G Beads were purchased from Thermo Scientific. The western chemiluminescent HRP substrate was purchased from Millipore Corporation.

### Western blotting

All cells were lysed in RIPA (Solarbio) supplemented with protease and phosphatase inhibitor cocktail (100×, Thermo Fisher) and lysed for 10 min on ice. The supernatant was mixed with 1/4 volume of 5× loading dye. The mixtures were then heated at 95°C and stored at –80°C. The samples were separated by SDS-PAGE and transferred onto nitrocellulose membranes. The membranes were then blocked with 5% nonfat milk (BD) in 1× Tris-buffered saline and 0.1% Triton 100 for 1 h while shaking at room temperature. Next, the membranes were incubated with primary antibodies and horseradish peroxidase–conjugated secondary antibodies. Bands were visualized using the Kodak film exposure detection system. The film was scanned, and the band intensity was analyzed using Quantity One software.

### Quantitative real-time PCR

Total RNA was extracted from cultured cells using TRIzol reagent (Invitrogen). cDNA was generated by reverse transcription with commercial PrimeScript RT Master Mix (Takara). Primer pairs (Table S12) were designed using Primer Premier Software 5.0 (Premier Biosoft International, Palo Alto, CA) and synthesized by Invitrogen. Quantitative real-time PCR was performed in triplicate wells of a 96-well reaction plate on an ABI 7500 PCR System (Applied Biosystems). GAPDH was used as the endogenous control. The 2^−ΔΔCt^ method was used to calculate expression relative to the internal control. The data were analyzed using ABI 7500 SDS software v.1.3.

### Luciferase assays

For IFNB1 transcriptional activity assays, 100 ng of pGL3-IFNB1 luciferase plasmid was cotransfected with 20 ng of pRL-TK vector into the cells using jetPRIME Transfection reagent (Polyplus). At 24 h after transfection, the cells were harvested according to the manufacturer’s protocol (Promega), and firefly and Renilla luciferase signals were measured using a dual luciferase reporter assay system (Promega) on a Promega GloMax 96 machine (Promega) according to the protocol provided by the manufacturer.

### Northern blotting

For northern blotting, total RNA was isolated from A549 cells using TRIzol reagent. Probes (Custom LNA mRNA Detection) were designed and synthesized by Exiqon and were also used for FISH. Northern blotting was performed as described previously (46).

### 5’ and 3’ RACE

The 5’ and 3’ RACE analyses were performed using the SMARTer RACE 5’/3’ Kit (Clontech) according to the manufacturer’s instructions. The RACE PCR products were cloned into pMD-19Tvector (Takara) and sequenced.

### RNA pull-down assays

RNA pull-downs were performed as described (46). *In vitro* biotin-labeled RNAs (lnc*AVAN* and its antisense RNA) were transcribed with biotin RNA labeling mix (Roche) and T7 RNA polymerase (Roche), treated with RNase-free DNase I (Promega) and purified using the RNeasy Mini Kit (QIAGEN). Biotinylated RNA was incubated with cell lysate, and precipitated proteins were separated via SDS-PAGE. Then the gel was stained using SilverQuest™ Silver Staining Kit (Invitrogen™) according to the manufacturer’s instructions. Following silver staining, specific bands in sense group and control bands with equal molecular weight in antisense group were cut and fed into nanoLC-LTQ-Orbitrap XL for mass spectrometry analysis.

### RNA fluorescence *in situ* hybridization (RNA-FISH)

RNA-FISH was performed as described previously(51). Hybridization was carried out using DNA probe sets (Biosearch Technologies) according to the protocol provided by Biosearch Technologies. Cells were observed on a FV1000 confocal laser microscope (Olympus).

### RNA immunoprecipitation (RIP) and chromatin immunoprecipitation (ChIP) assays

RIP assays were performed as described (51) with minor modification. Briefly, after incubation, the magnetic beads were washed with high salt lysis buffer (containing 500mM NaCl) 5 times. RIP products were analyzed by qRT-PCR using the primer pairs listed in Table S12. ChIP was performed as described (67). The ChIP-enriched FOXO3a promoter was quantified by qPCR using the primer pairs listed in Table S12.

### ChIRP

ChIRP was performed according to Chu *et al* (68). Briefly, seven non-overlapping antisense DNA probes targeting *AVAN* were designed (http://www.singlemoleculefish.com), and eight probes targeting LacZ were also designed as non-specific controls. All probes were biotinylated at the 3’ end with an 18-carbon spacer arm (Invitrogen). 20 million cells were cross-linked by 1% formaldehyde. Cross-linked cells were lysed in lysis buffer (50 mM Tris pH 7.0, 10 mM EDTA, 1% SDS, add DTT, PMSF, protease inhibitor, and RNase inhibitor before use) at 100mg/ml on ice for 10 min, and sonicated using Ultrasonic Cell Crusher on ice until the bulk of DNA smear is 100-500bp. The cell lysate was separated into two equal aliquots, one for hybridizing with probes targeting *AVAN*, and the other for probes targeting LacZ as control. Next, C-1 magnetic beads were added to the probe-chromatin mixture. After five total washes with washing buffer (2×SSC, 0.5% SDS, add DTT and PMSF before use), the beads were separated into two parts, 1/10 for RNA elution and 9/10 for DNA elution or protein elution. For elution of the RNA, the beads were treated with Protease K Buffer and RNA was extracted with Trizol reagent. For elution of the DNA, beads were treated with DNA Elution Buffer (50mM NaHCO_3_, 1% SDS, add 100ug/ml RNase A and 100U/ml RNase H before use), and then DNA was isolated via phenol:chloroform:isoamyl alcohol extraction and subjected to qPCR. For elution of the protein, beads were treated with β-mercaptoethanol at a final concentration of 2.5% in 1× NuPAGE LDS Sample Buffer (Life Technologies, NP0007) at 96 °C for 30 min.

### Statistical analyses

All data are presented as the means ± SEM. Data were analyzed using Student’s t test. Survival data were analyzed by Kaplan-Meier survival analysis, and single time-points were analyzed using ANOVA. All analyses are performed using Instat software (Version 5.0, GraphPad prism). p<0.05 was considered statistically significant.

## Acknowledgments

This work was supported in part by funding from the National Programs for High Technology Research and Development of China (SS2015AA020924), the National Major Research and Development Program of China (No. 2017YFC1200800) and the Natural Science Foundation of China (81771700).

## Author contributions

Authors P.Y., R.C. and X.W. conceived and designed the experiments; C.L., L.L., Q.L., S.C., K.W., L.Z., M.X. and H.G. performed the experiments; Y.D., C.W., Z.Z., L.Z., J. S., Z. L. and J.L. analyzed the data; and C.L., L.L., K.W. and Q.L. wrote the manuscript.

