## Supplementary figures and images for "Long noncoding RNA *AVAN* promotes antiviral innate immunity by interacting with TRIM25 and enhancing the transcription of FOXO3a"

### Supplemental Figure 1

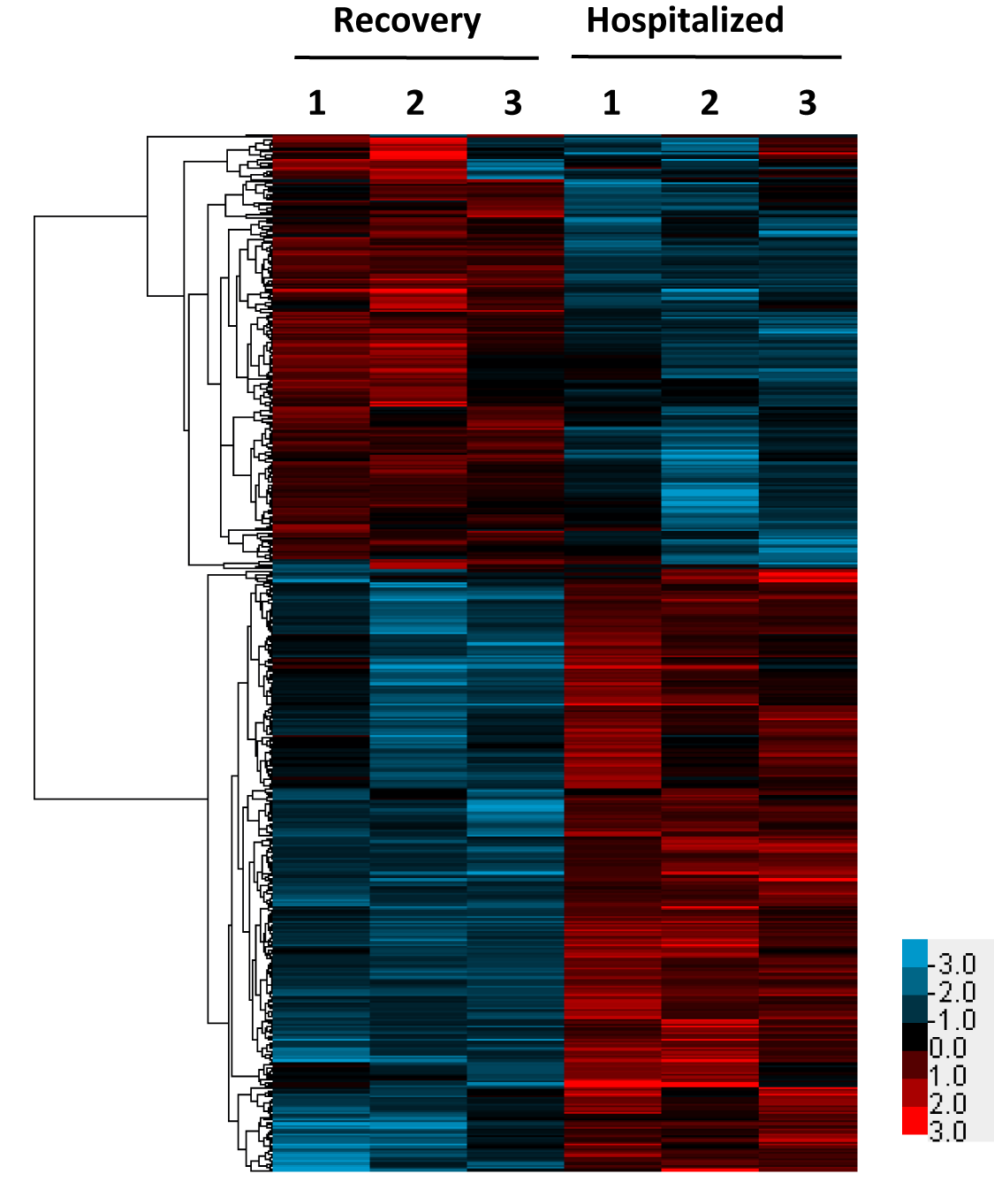

### Supplemental Figure 2

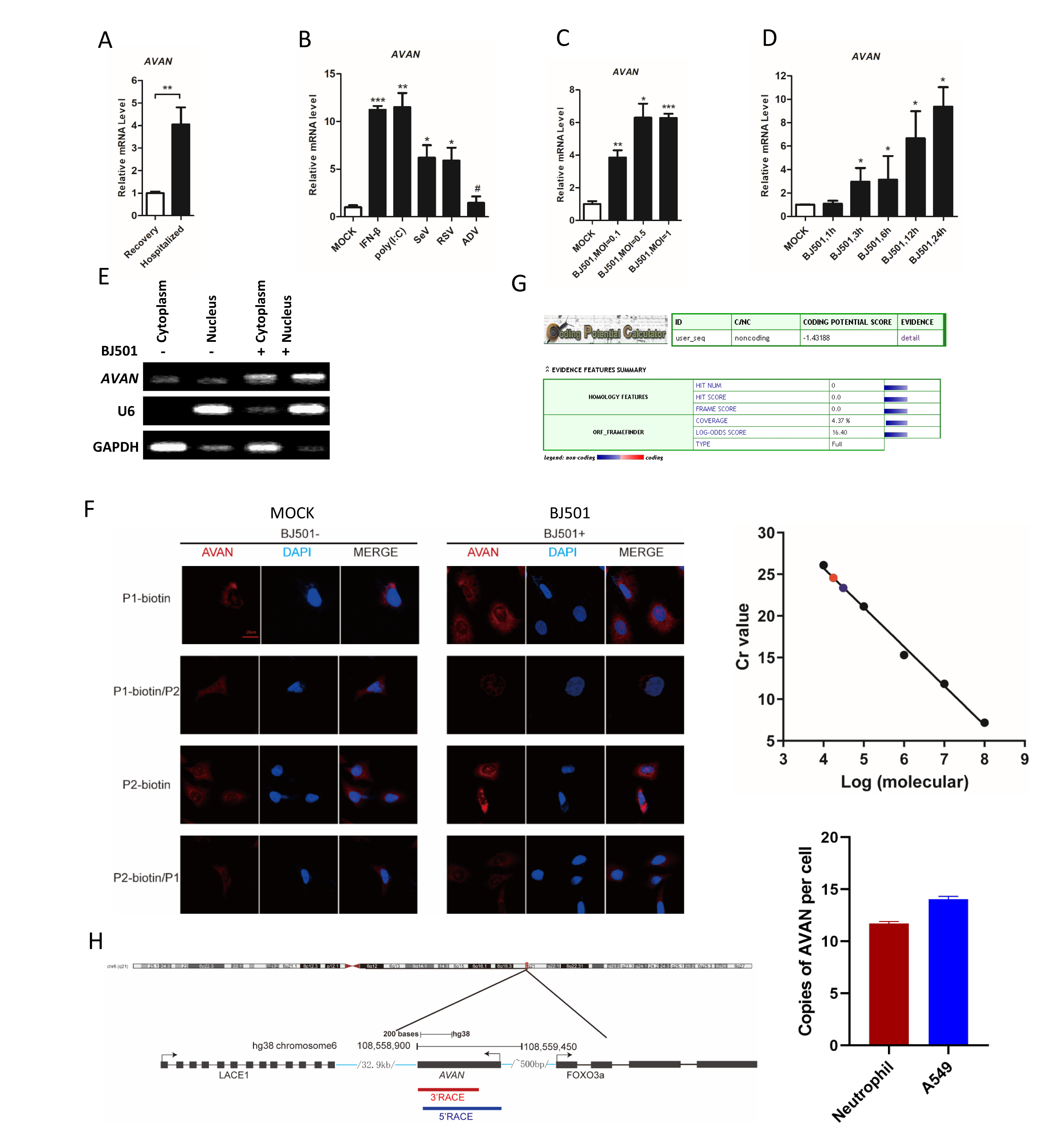

### Supplemental Figure 3

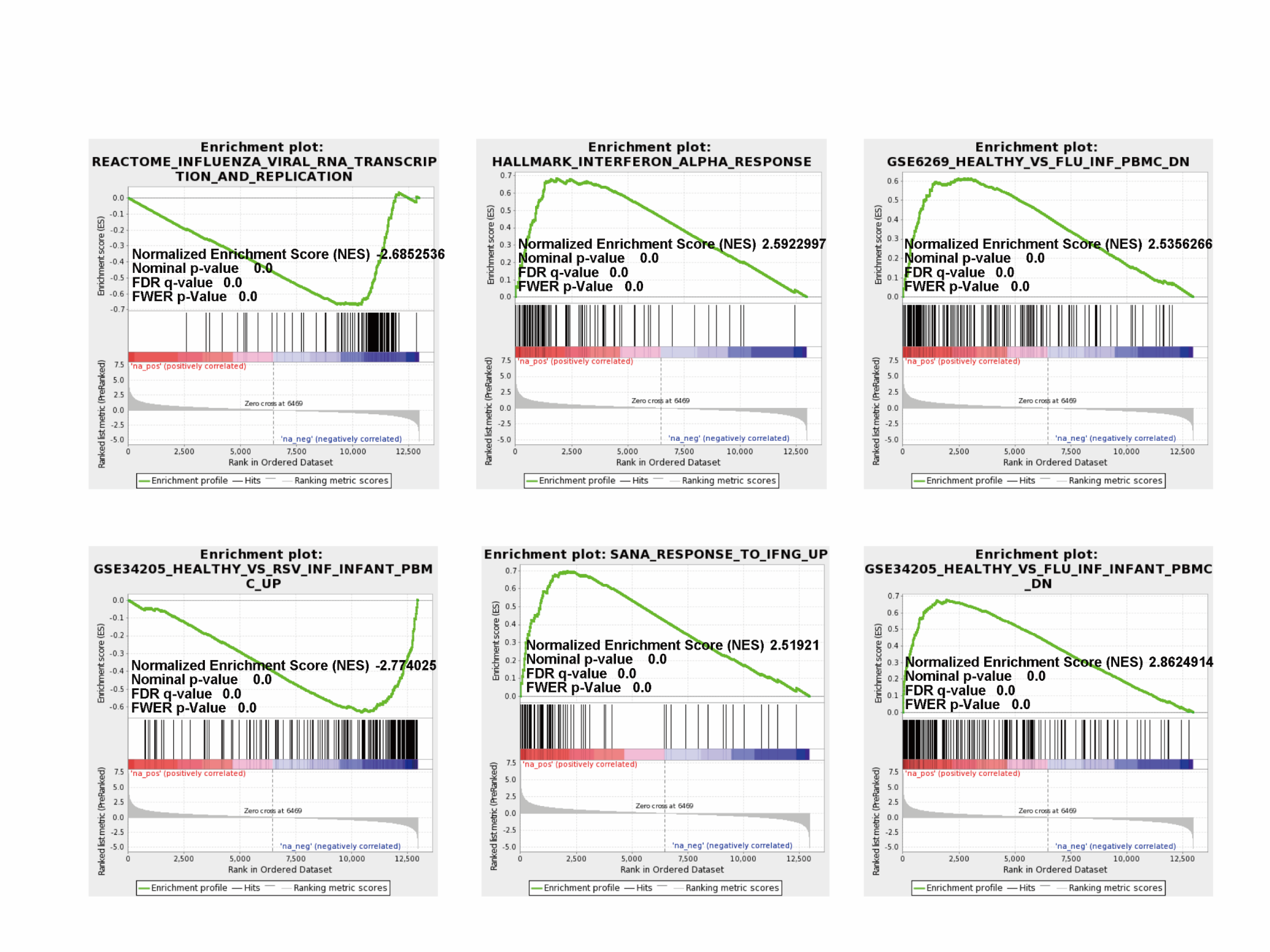

### Supplemental Figure 4

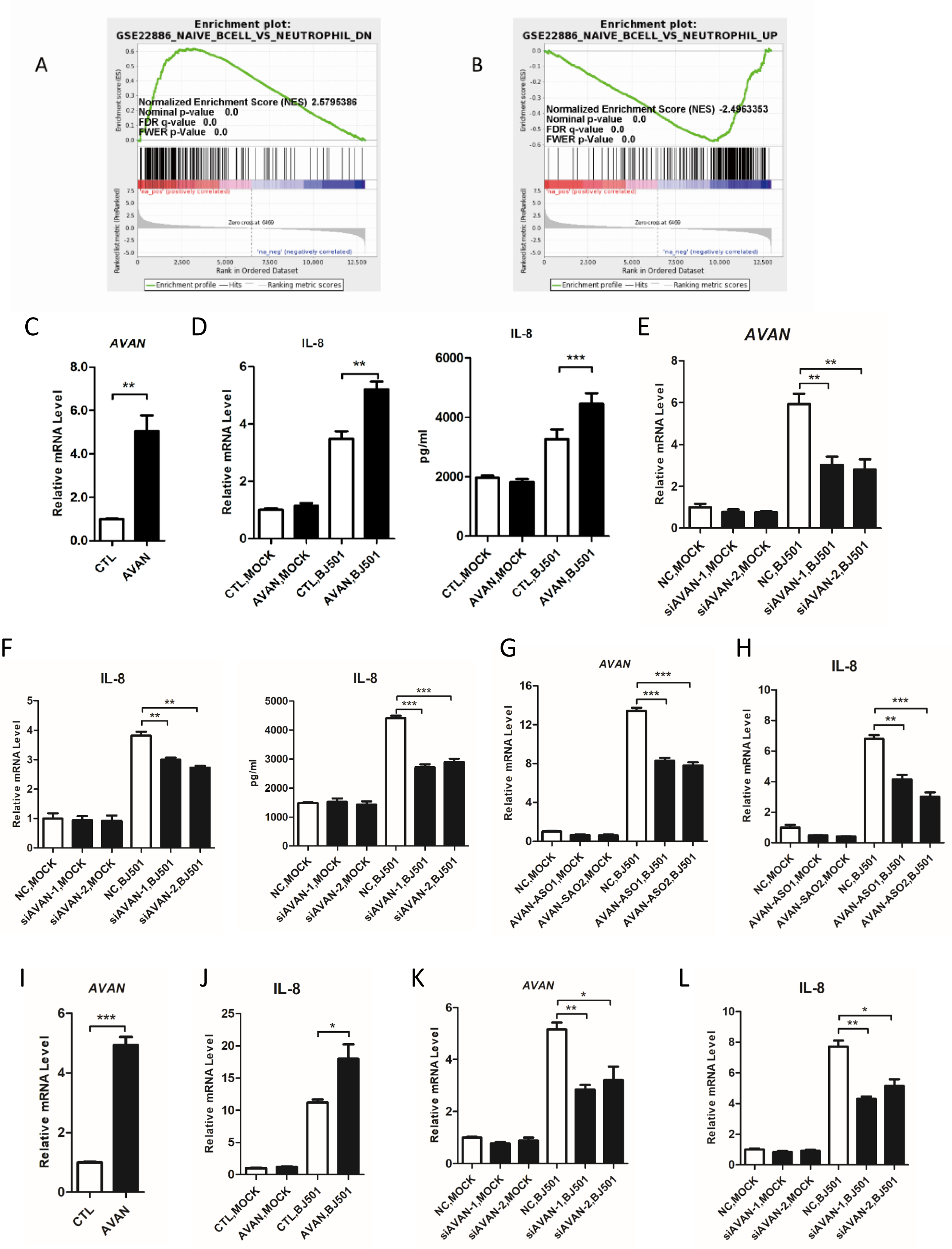

### Supplemental Figure 5

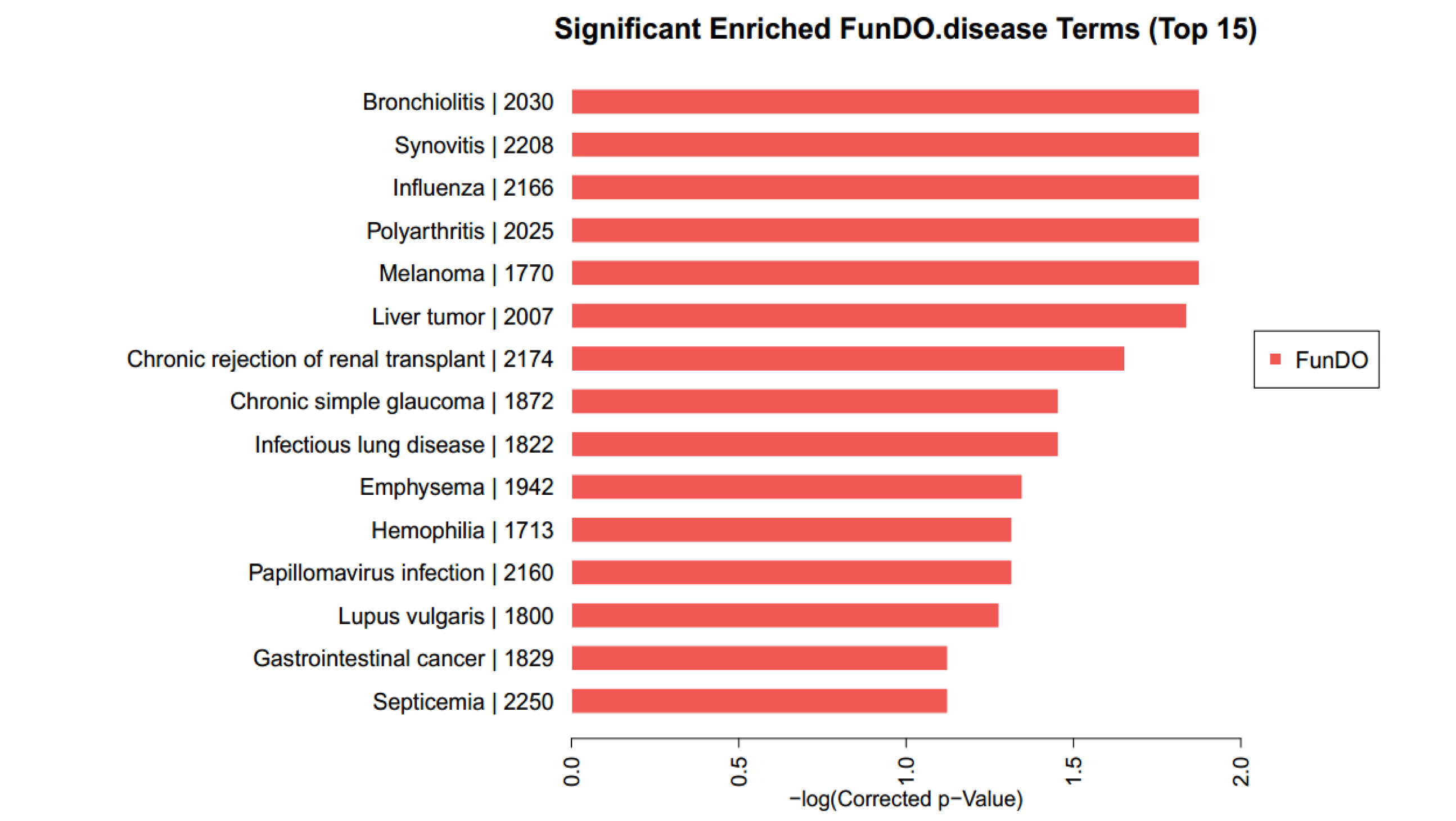

### Supplemental Figure 6

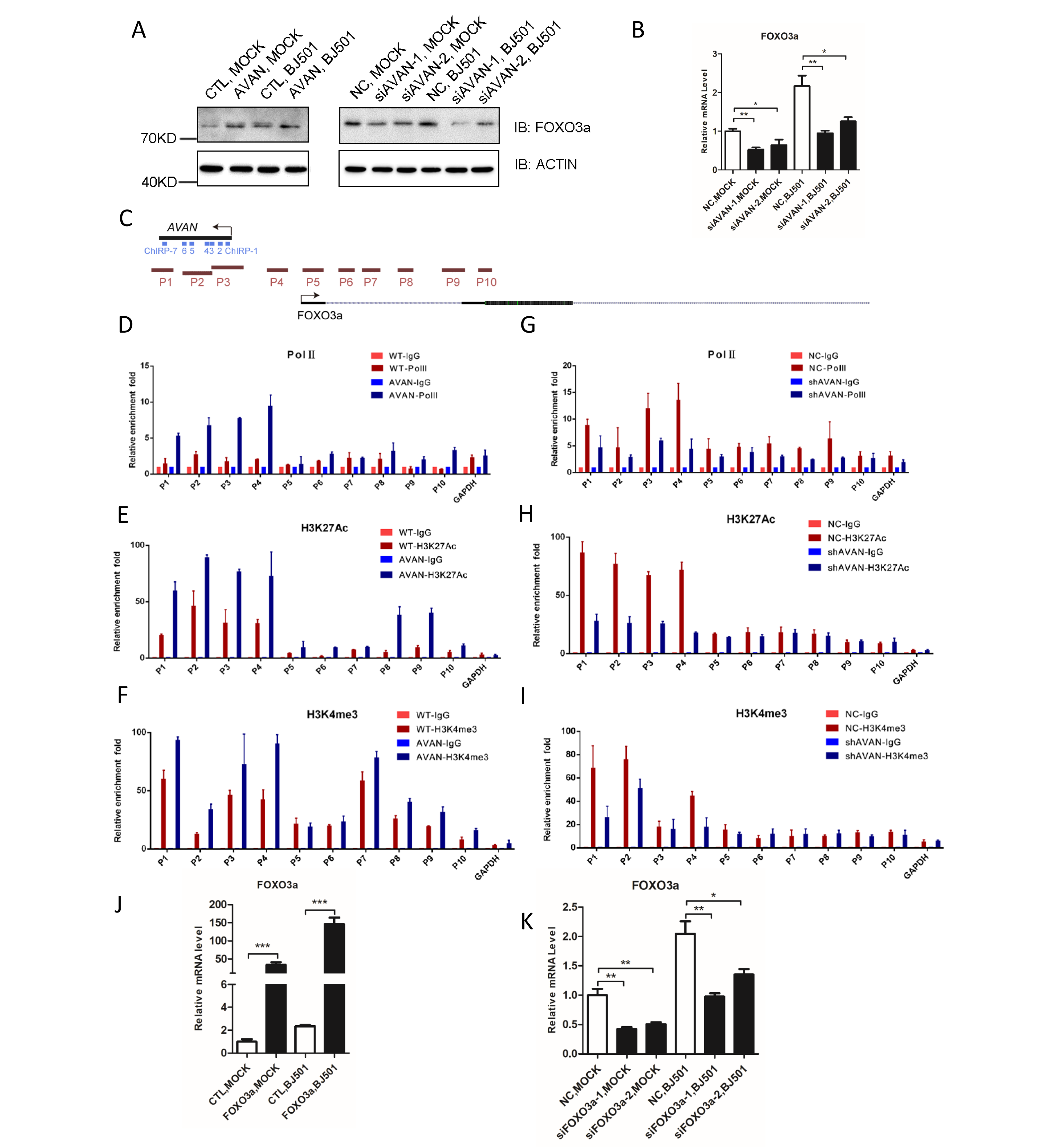

### Supplemental Figure 7

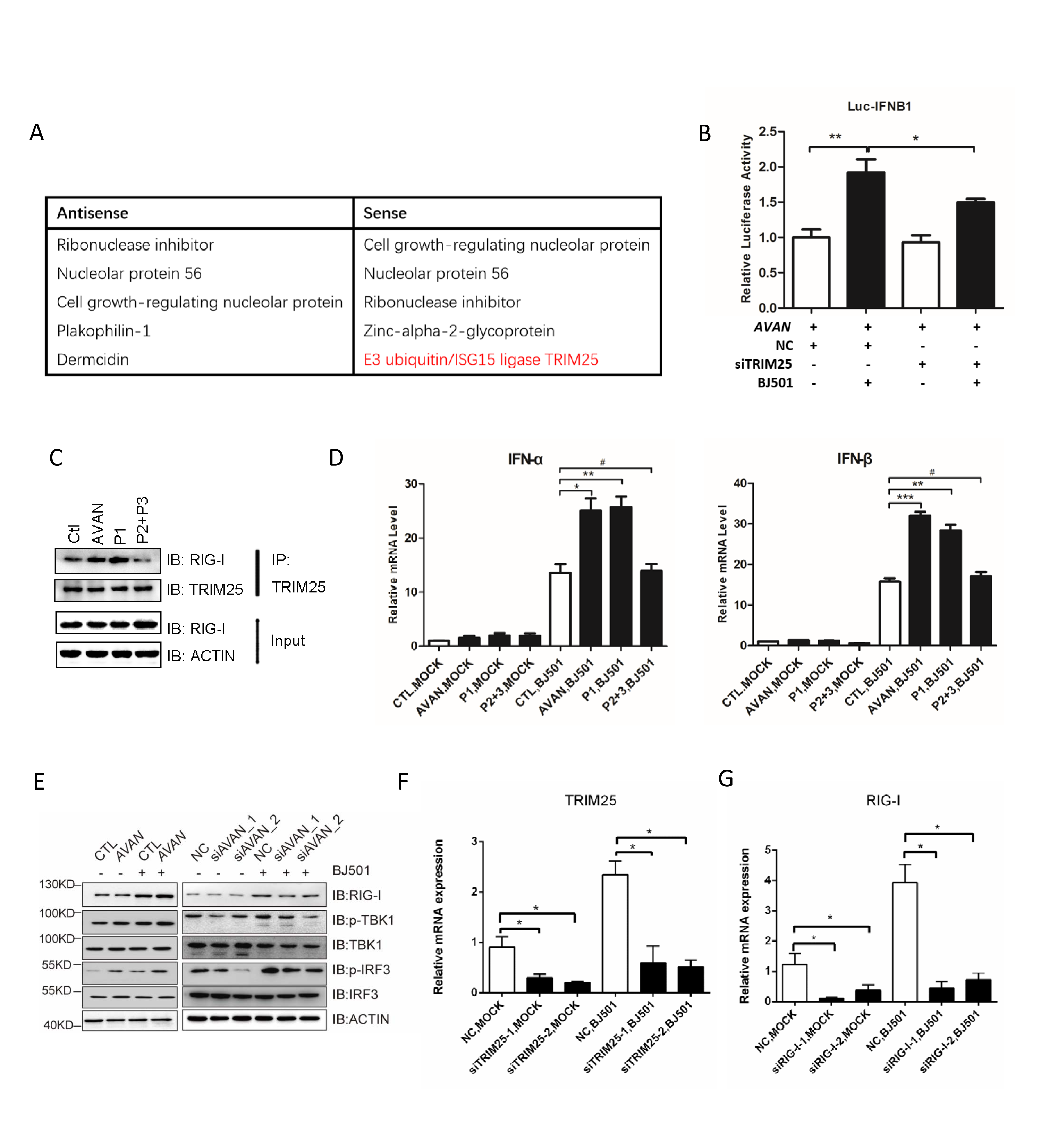

### Supplemental Figure 8

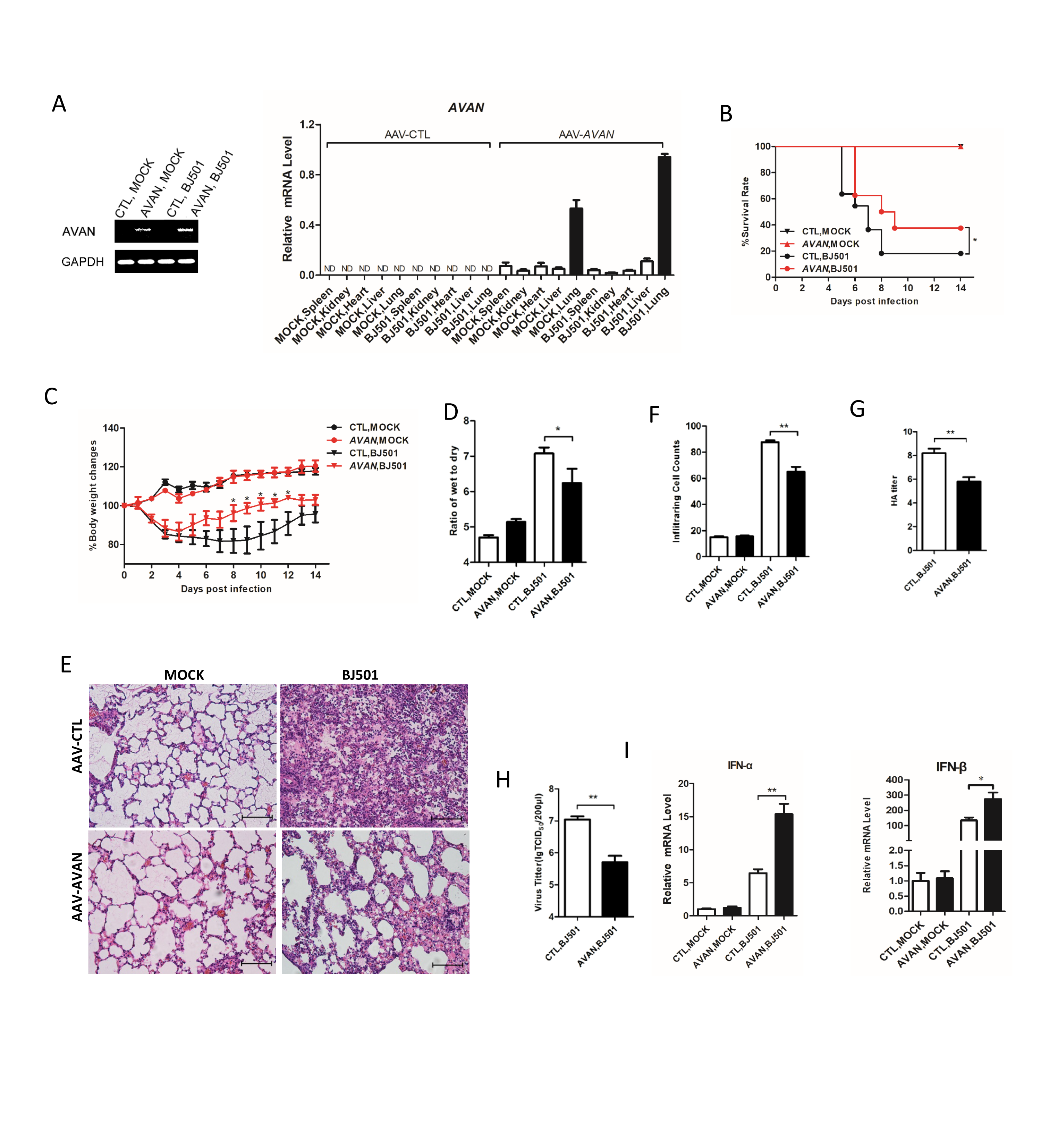
