## Supplemental Table 1 for "Long noncoding RNA *AVAN* promotes antiviral innate immunity by interacting with TRIM25 and enhancing the transcription of FOXO3a"

| Table S1. basic information of patient recruited | |
| --- | --- |
| Characteristic, Underlying Conditions and Outcomes of 63 Patients Infected with 2009 Influenza A(H1N1) Virus in China | |
| Characteristic |  |
| Male sex-no. (%) | 36(55.56) |
| Age-yr |  |
| Mean | 41.47 |
| Range | 1-89 |
| Age group-no. (%) |  |
| <19 | 7(11.11) |
| 20-25 | 11(17.46) |
| 26-30 | 8(12.69) |
| 30-35 | 13(20.63) |
| >35 | 24(38.1) |
| Coexisting conditions-no. (%) |  |
| Hypertension | 7(11.11) |
| Chronic obstructive pulmonary disease | 5(7.93) |
| Asthma | 3(4.76) |
| Diabetes | 1(1.59) |
| Cancer | 4(6.35) |
| Coronary heart disease | 6(9.52) |
| Diagnosis-no. (%) |  |
| Mild | 17(26.98) |
| Hospitalized but not critical | 30(47.62) |
| Critical | 16(25.4) |
| Lung injury extent-no. (%)* |  |
| No lung injury | 22(34.92) |
| Acute lung injury | 32(50.79) |
| Acute respiratory distress syndrome | 9(14.29) |
| Outcomes-no. (%) |  |
| Recovery | 55(87.3) |
| death | 8(12.7) |
