## Supplemental Table 2 for "Long noncoding RNA *AVAN* promotes antiviral innate immunity by interacting with TRIM25 and enhancing the transcription of FOXO3a"

**Table S2. LncRNA-AVAN sequence from RACE**

>hg38_dna range=chr6:108558901-108559417 AAGGATACGTCTTTTTTTAAAAGTCAAAAATGCCCACGTAGCGGTTTTGTCGTCATTTACAAATTAGATTAAAATTATTTCCCCTTCTCAGTTGTTCGGAATCTGCTCTTGAATGTAGGACGGGGGTAAATTTGCATTTCTTCCGGAGTTAGTTACACCTCTCCCCCGTTTGCCATGAAACCCATGAAAAGTGAAACACGACTGTGGTCCACCGTCCGAGGTGAGAGACTCAGTGCAAACCTTTTGGTGCCTGATCGAATCTGTGAGCATCAGCTTATCGAAGTTGTGCAGGCACCAACATCTTCGCCGTTCAGTATTTCCACACCGCGATAACCCTGCCCATCTTGCGGCCAGCATCTTATCTCGGGTGTTACCTGGTGTCGCGTTCTAACAGGGAACCGGACACCGGGGCTGGCCCAGAGCCCGGCGGGCGGACGCGCAGGCCAGGGACCCCCTGGTACCAGGCGCCCTGCCCGGAGTCAGGGAGGCCACGCCGCTGCGCTCCTTGGAGCGTCGG
