## Supplemental Figure legend for "Long noncoding RNA *AVAN* promotes antiviral innate immunity by interacting with TRIM25 and enhancing the transcription of FOXO3a"

**Figure S1 Altered mRNA expression in neutrophils during IAV infection.**

Cluster heat map showing altered mRNA expression in IAV-infected patient neutrophils compared with recovery-stage samples from RNA-seq data (FC > 2; p<0.05).

**Figure S2 Time- and dose-dependent AVAN activation during IAV infection and characterization of AVAN.**

(A) AVAN up-regulation in IAV-H7N9-infected patient (n=3; means ± SEM; **p<0.01).

(B) AVAN expression in A549 cells treated with IFN-β (10ng/ml), poly I:C (0.5μg/ml), sendai virus (SeV) (MOI=1), respiratory syncytial virus (RSV) (MOI=1) or adenovirus (ADV) (MOI=1) for 24 h by qRT-PCR analysis (n=3; means ± SEM; **p<0.01; ***p<0.001).

(C and D) A549 cells were infected with BJ501 at MOI=1 for the indicated time or at the indicated MOI for 24 h. qRT-PCR was performed to measure AVAN expression (n=3; means ± SEM; **p<0.01).

(E) Fractionation of BJ501-infected A549 cells followed by RT-PCR analysis (MOI=1) at 24h post-infection. U6 RNA served as a positive control for nuclear gene expression, and GAPDH served as a positive control for cytoplasmic gene expression.

(F) *AVAN* intracellular localization visualized by RNA-FISH in A549 cells stimulated or not with BJ501 (MOI=1) for 24 h. DAPI, 4’,6-diamidino-2-phenylindole. Probe 1-biotin and probe2-biotin, biotinylation AVAN probes; probe1 and 2, no labeled free AVAN probes.

(G) The coding potential of AVAN according to the Coding Potential Calculator (http:// cpc.cbi.pku.edu.cn/programs/run_cpc.jsp).

(H) Schematic diagram showing full-length AVAN gene locus and RACE products.

**Figure S3 GSEA analysis of related gene co-expression networks from the sequencing data.**

**Figure S4 Altered AVAN expression affects chemokine expression levels.**

Cells were transfected with AVAN or negative control and AVAN siRNAs, ASOs or negative control siRNAs, negative control ASO, as indicated.

(A and B) GSEA data and GO analysis of genes related to AVAN.

(C and E) The efﬁciency of AVAN overexpression (C) and siRNA-based knockdown (E) was determined by qRT-PCR in HL60 cells (n=3; means ± SEM; **p<0.01;).

(D and F) IL-8 mRNA expression at 12 h post-infection determined by qRT-PCR and IL-8 levels in cell culture supernatants determined by ELISA in HL60 cells (n=3; means ± SEM; **p<0.01; ***p<0.001).

(G) The efﬁciency of ASO-based knockdown of AVAN was determined by qRT-PCR in A549 cells (n=3; means ± SEM; **p<0.01;).

(H) IL-8 mRNA expression at 24 h post-infection determined by qRT-PCR in A549 cells (n=3; means ± SEM; **p<0.01; ***p<0.001).

(I and K) The efﬁciency of AVAN overexpression (I) and siRNA-based knockdown (K) was determined by qRT-PCR in primary human neutrophils (n=3; means ± SEM; *p<0.05; **p<0.01; ***p<0.001).

(J and L) IL-8 mRNA expression at 12 h post-infection determined by qRT-PCR in primary human neutrophils (n=3; means ± SEM; *p<0.05; **p<0.01).

**Figure S5 FunDO diseases constructed from differentially expressed mRNAs in response to AVAN with the listed score; the top 15 are presented.**

**Figure S6 AVAN enhances FOXO3a expression *in cis***

Cells were transfected with AVAN or negative control and AVAN siRNAs or negative control siRNAs, as indicated.

(A) Western blotting of FOXO3a protein expression in BJ501-infected (MOI=1) at 24h post-infection A549 cells after overexpressing or knocking down AVAN.

(B) FOXO3a expression in BJ501-uninfected (left) and -infected (right) HL60 cells were measured by qRT-PCR after AVAN knockdown (n=3; means ± SEM; *p<0.05; **p<0.01).

(C) Schematic diagram showing the probes and primers used in ChIRP and ChIP.

(D) H3K4me3 and H3K27ac levels and Pol II binding of the FOXO3a promoter were analyzed through ChIP followed by qPCR in A549 cells after overexpressing (D-F) or knocking down (G-I) AVAN.

(J and K) The efﬁciency of FOXO3a overexpression (J) and knockdown (K) was determined by qRT-PCR in A549 cells that were infected by BJ501 or not (MOI=1) at 24h post-infection (n=3; means ± SEM; *p<0.05; **p<0.01; ***p<0.001).

**Figure S7 AVAN direct binds to TRIM25 and enhances the antivirus immune response.**

(A) The list of proteins identified by MS in *AVAN* sense and antisense group, and particularly TRIM25 was pulled down by AVAN in IAV-infected A549 cells but not by the AVAN antisense.

(B) AVAN was overexpressed in all four groups, meanwhile endogenous TRIM25 was knocked down in HEK-293T cells and then treated with IAV-BJ501 or not. IFNB1 transcription measured using dual luciferase reporter assays in HEK-293T cells (n=3; means ± SEM; *p<0.05; **p<0.01).

(C) TRIM25 co-immunoprecipitation with proteins from lysates of BJ501-infected A549 cells transfected with different AVAN fragments (MOI=1) at 24h post-infection, followed by immunoblotting. Anti-TRIM25 and anti-RIG-I antibodies were used for immunoprecipitated.

(D) IFN-alpha (left) and IFN-beta (right) expression analyzed by qRT_PCR upon *AVAN* or *AVAN* truncates (Fig 6G upper panel) transfection in A549 cells that were infected by BJ501 or not (MOI=1) at 24h post-infection.

(E) Western blot analysis of RIG-I signaling in A549 cells transfected with *AVAN* or siRNAs.

(F and G) The efﬁciency of knocking down TRIM25 (F) and RIG-I (G) was determined by qRT-PCR in A549 cells that were infected by BJ501 or not (MOI=1) at 24h post-infection.

**Fig S8 *AVAN* protects mice from virus infection**

(A left) *AVAN* expression in the lung of AAV2/9-*AVAN*-treated or control mice measured by RT-PCR.

(A right) *AVAN* expression in the lung, spleen, liver, kidney, heart of AAV2/9-*AVAN*-treated or control mice measured by qRT-PCR.

(B and C) Four-week-old wild-type B6 mice were inoculated with 10^5.125^ TCID_50_ BJ501 virus. Survival rates (B) and body weight changes (C) of wild-type mice (n=10 for each group) were monitored for 2 weeks after BJ501 challenge (*p<0.05).

(D) Wet:dry ratios of lung tissues (n=6 for each group) at 5 DPI (*p<0.05).

(E and F) HE-stained images (E) and infiltrating cell counts (F) (n=100 fields) in lung tissues at 5 DPI (magnification=200×), (**p<0.01).

(G and H) Lung HA titer (G) and virus titer (H) at 5 DPI (**p<0.01).

(I) IFN-α and IFN-β expression in the lung of mice treated with AAV2/9-*AVAN* or control vector measured by qRT-PCR (n=3; means ± SEM; *p<0.05; **p<0.01).
